# Nonadjacent Dependencies and Syntactic Structure of Chimpanzee Action During a Natural Tool-Use Task

**DOI:** 10.1101/2024.03.25.586385

**Authors:** Elliot Howard-Spink, Misato Hayashi, Tetsuro Matsuzawa, Daniel Schofield, Thibaud Gruber, Dora Biro

## Abstract

The hierarchical organization of sequential behaviour, and the ability to compensate for nonadjacent sequential dependencies, are fundamental and interrelated abilities supporting complex human behaviours, including language and tool use. To understand how the cognition underlying these structural properties of human behaviour evolved, we can gain valuable insight from studying the sequential behaviours of nonhuman animals. Among the behaviours of nonhuman apes, tool use has been hypothesised to be a domain of behaviour which likely involves hierarchical organization. However thus far, evidence supporting this hypothesis comes from methodologies which have been criticised in their objectivity. Additionally, the extent to which nonadjacent dependencies appear in primate action sequences during tool use has not been formally investigated. We used estimations of mutual information (MI) – a measure of dependency strength between sequence elements – to evaluate both the extent to which wild chimpanzees produce nonadjacent dependencies during a naturalistic tool-use task (nut cracking), as well as how sequences of actions are organized during tool use. Half of adult chimpanzees produce nonadjacent dependencies at significantly greater sequential distances than comparable, nonhierarchical Markov models, including when repeated actions had been accounted for. Additionally, for the majority of chimpanzees, MI decay with increasing sequential distance included a power-law relationship, which is a key indicator that most chimpanzees draw upon forms of hierarchical structuring when organizing behaviours during tool use. Our analysis offered the greatest support for a system of organization which involved the production of short subroutines of actions (2-8 actions), which are hierarchically arranged into sequences – a finding which is consistent with previous qualitative descriptions of ape tool-use behaviours. Interindividual variability was detected within our analysis in both the distance dependencies were detected, and the most likely structuring mechanism for sequential action organization. We discuss these results in light of possible interindividual variation, in addition to methodological considerations for applications of MI estimations to sequential behaviours. Moreover, we discuss our main findings alongside hypotheses for the coevolution of complex syntax in language and tool-action across hominin evolutionary history.

## 1. Introduction

Many behaviours of both humans and non-human animals (henceforth animals) can be considered sequences of discrete actions. Therefore, understanding how both humans and animals generate, store and retrieve such sequences is key to understanding the cognition supporting these serial behaviours. Humans in particular exhibit a number of sequential behaviours which are argued to be uniquely complex in comparison to the analogous behaviours of non-human animals, including language (Berwick and Chomsky, 2015; Bolhuis et al., 2014; Hauser et al., 2014, 2002); the production and appreciation of music (Fitch, 2013; Fitch and Martins, 2014); and the manufacture and use of highly sophisticated tools (Davidson and McGrew, 2005; Greenfield, 1991; McGrew, 1993; Vaesen, 2012). These elaborate sequential behaviours all share similar mechanisms of organization, where the atomic units of each behaviour (words, musical notes, and manual actions) are organized hierarchically within the sequence (Asano and Boeckx, 2015; Chomsky, 2002, 1956; Fitch and Martins, 2014; Greenfield, 1991; Lerdahl and Jackendoff, 1983; Stout et al., 2021). Hierarchical sequential organization involves the association of elements into short subroutines, which are themselves organized to produce wider strings of behaviour. In language, this can be seen through the organization of words into clauses, clauses into sentences, and sentences into longer structures (such as the paragraphs you are reading now; Chomsky, 2002; Coopmans et al., 2021; Fitch and Martins, 2014). Similarly in tool-use behaviours, actors parse the wider goal of their behaviour into subgoals, which themselves may be recursively split into smaller nested subgoals. Actors then generate sequential behaviours for tool use by generating actions which address the nested subgoals in turn (Byrne et al., 2013; Coopmans et al., 2021; Pastra and Aloimonos, 2012). For example when making a cup of tea, an actor may partition this behaviour into the subgoals of: *fill the kettle with water*, *boil the kettle*, and *brew the tea*, before parsing their first subgoal (*fill the kettle with water*) into a number of smaller goals including *open the kettle*, *turn on the tap*, *hold the kettle below the water stream*, *turn off the tap* and *close the kettle lid*. This process can be traced until one reaches the level of individual manual actions (for example, to *turn off the tap*: grasp the handle, twist, and release).

The hierarchical organization of human language, music, and actions during tool use results in these domains of behaviour sharing a distinctive property: the emergence of nonadjacent dependencies between sequence elements. Nonadjacent dependencies can emerge during hierarchical organization through the embedding of subroutines within other subroutines. Within language, this is reflected when a clause is embedded within another clause (Chomsky, 2002, 1956; Fitch and Martins, 2014; Jäger and Rogers, 2012; Lashley, 1951; Sainburg et al., 2022, 2019); e.g. in the sentence “*The weather is horrible!*”, a prepositional phrase may be embedded to produce the sentence “*The weather [outside the window] is horrible!”*. Such a sentence may then feature nonadjacent dependencies, where the verb ‘is’ and adjective ‘horrible’ both relate to the noun at the head of the phrase (the weather), despite being closer to another noun (the window) in linear space and/or time. Another example from language includes subject-verb agreements, where the nature of the subject (gender, number, etc.) may influence the morphology of its corresponding verb, irrespective of how far separated they are in linear space and/or time (Mel’čuk, 1988; Rispens and Soto de Amesti, 2017). Likewise during sequential actions, nonadjacent dependencies occur when a series of actions addressing one subgoal are embedded within a sequence of actions addressing another goal. For example, when opening a door an actor may grasp, turn, and push the handle. However, if the door is locked, the actor may grasp the handle, then begin eliciting a number of actions to open the lock (thus completing the embedded subgoal: undo the lock), before continuing to turn and push the handle to open the door. In such an example, grasping a handle always precedes turning a handle; however, they may be separated by an embedded string of actions, producing a nonadjacent dependency similar to what is seen in language.

The fact that hierarchical organization, and its resultant nonadjacent dependencies, are prolific across a number of complex and open-ended human behaviours suggests that hierarchical organization is necessary for the evolution of behaviours which are unbounded in their generativity, but maintain the ability to be employed in a directed manner (for further discussions surrounding this trade-off see Byrne et al., 2013; Chomsky, 2002; Dawkins, 1976; and Isac and Reiss, 2013). However, hierarchical sequential organization entails cognitive costs. This is particularly true for working memory, since hierarchical organization requires an individual to remember, trace and match dependencies between elements across the levels of a hierarchy whilst behaviours are performed in real time (Fitch and Hauser, 2004; Wilson et al., 2020). It is therefore reasonable to expect highly hierarchical behaviours (i.e. those which span many nested levels of organization) to evolve when the benefits of high behavioural flexibility outweigh the costs of maintaining the underlying neural architecture for more sophisticated cognition. This has led to an explosion of research in recent decades, aiming to address how and why humans evolved the cognition to support the hierarchical organization across such an array of behavioural domains (Asano and Boeckx, 2015; Coopmans et al., 2021; Fitch and Hauser, 2004; Fitch and Martins, 2014; Friederici, 2020; Greenfield, 1991; Hauser et al., 2002; Kemmerer, 2022; Lerdahl, 2015; Mielke and Carvalho, 2022).

Comparative research into the sequential behaviours of nonhuman animals may offer key insights into the evolution of hierarchical behavioural organization, particularly through the investigation of behaviours which are analogous to human communication, music and tool use (Berwick et al., 2012; Fitch and Hauser, 2004; Hauser et al., 2002; Hayashi, 2015; Matsuzawa, 1996; Sainburg et al., 2019, 2019). Whilst hierarchical organization in human behaviour has been theorized, and empirically investigated, since the mid-20^th^ century, the idea that animals may also possess the ability to organize behaviours hierarchically emerged more slowly, possibly due to the widely held belief that animals lacked the behavioural sophistication (and underlying cognition) for hierarchical behaviours (Terrace, 2005). Instead, animal sequential behaviours were assumed to be produced through non-hierarchical reflexive chains: when an action or call was externalised, it would trigger the externalization of the next action or call, and so on to produce an extended behavioural sequence (Hauser et al., 2002; ten Cate and Okanoya, 2012; Terrace, 2005). The rules which animals employed to create such behavioural sequences were therefore theorized to be extremely simple, requiring no knowledge of the calls or actions they produced other than the one produced most recently (thus, highly reducing the burden on working memory due to the absence of nonadjacent dependencies; Wilson et al., 2020). Such reflexive chains can be replicated using Markov models: network models where each node represents a different possible element in a sequence, and the edges between nodes are weighted based on the likelihood a sequence will contain an adjacent transition between element types (Gagniuc, 2017).

Despite the historic pervasiveness of chaining models for explaining the sequential structure of animal behaviours, mounting evidence suggests that a number of different animal species employ hierarchical sequencing to produce behaviours in the wild. For example, hierarchical organization is found the songs of whales (Miksis-Olds et al., 2008; Suzuki et al., 2006) and birds (Sainburg et al., 2019), as well as in the behaviours of several primate species, including grouped hierarchical perceptions of social relationships in baboons (Bergman et al., 2003); the nested temporal organization of vocal elements during orangutan long calls (Lameira et al., 2024); and the mental organization of social games in chimpanzees (Mielke and Carvalho, 2022). Nevertheless, hierarchical organization does not yet appear to be a universal aspect of animal behaviours. For instance, many species appear to lack hierarchical organization in their communicative behaviours (Fitch et al., 2005; Hauser et al., 2002; ten Cate and Okanoya, 2012), suggesting that this phenomenon may be limited to species that employ complex broadcast signals. The extent to which animals employ hierarchical organization may also vary between behaviours, including for humans. For example in human language, whilst words are hierarchically organized based on the rules of a language’s syntax, the relationships between sound units within words (phonemes) can be characterized using their neighbouring associations, implying that hierarchical organization is not involved during the combination of phonemes into meaningful units (Kaplan and Kay, 1994). Further research is therefore required to clarify precisely which species of animals employ hierarchical organization in sequential behaviours, as well as the specific behaviours that exhibit hierarchical organization. This includes research into the behaviours of species which are phylogenetically proximate to humans - such as chimpanzees, bonobos, and other great apes - for whom hierarchical behaviours, if present, would have the highest likelihood of sharing homologous origins with similar behaviours of humans (Gruber and Clay, 2016).

Among the behaviours of wild chimpanzees – one of the most closely related great ape species to humans – tool use has been identified as a domain of behaviour which likely possesses high hierarchical complexity (Byrne et al., 2013; Byrne and Russon, 1998; Gontier, 2024). Previous descriptions of chimpanzee tool-use behaviours have included the possibility that some particularly complex forms of tool use require chimpanzees to understand hierarchical relationships between objects (Hayashi, 2007; Hayashi and Takeshita, 2022; Matsuzawa, 1996, 1991; Sanz and Morgan, 2010). Moreover, when considering how chimpanzees organize their behavioural sequences for tool use, previous models used qualitative descriptions of the stages of producing, using, and disposing of tools, over many hours of human observation (the chaîne opératoire; Boesch et al., 2020; Byrne and Russon, 1998; Carvalho et al., 2008; Estienne et al., 2017; Sanz and Morgan, 2010; Tonooka, 2001). These observations have been used to suggest that, when engaging in tool-use behaviours, chimpanzees (and other apes) generate behavioural sequences hierarchically, through the decomposition of tool-use behaviours into a series of nested subgoals, which are themselves addressed through the generation of short subroutines of manual actions (Byrne et al., 2013; Byrne and Russon, 1998). Hierarchical organization is theorized to offer an adaptive advantage for successful tool-use behaviours in comparison to associative chaining, as organizing sequences hierarchically would allow for apes to flexibly reorganize subgoals of behaviours based on the local context of their environments, rather than repeatedly following a ‘fixed action pattern’ (Byrne et al., 2013). For example when organizing actions hierarchically, an individual may omit optional steps (a chimpanzee may not need to puncture a hole in a termite mound if one already exists; Byrne et al., 2013), or through embedding extra steps within a behavioural sequence (e.g. an individual may interrupt a behavioural sequence to adjust or swap over tool items, which are not necessary for the behavioural sequence, but may aid an individual in manipulating the tools during in a particular instance).

Despite chimpanzee tool use possessing many signs of hierarchical organization, the majority of evidence for hierarchical organization in chimpanzee tool use comes from qualitative descriptions of behaviours. Descriptive arguments for behavioural organization have been pointed out on numerous occasions to be at risk of subjective bias, as even though sequences can be described hierarchically, this does not stop them from being generated using computationally simpler mechanisms of sequence generation (Dawkins, 1976; Girard-Buttoz et al., 2022; Vereijken and Whiting, 1998). This argument is reinforced by the observation that humans are biased to perceive hierarchical relationships in sequences which lack any form of hierarchical organization (Ferrigno et al., 2020; Fitch, 2014), which in this instance could include the observation of animals’ behavioural sequences in the wild. There is consequently the need for new methodologies which are able to identify hierarchical properties of sequences objectively. Additionally, methodologies which also quantify the extent to which nonadjacent dependencies emerge between actions during such behavioural sequences would be particularly useful, as this is a property of chimpanzee tool use which has not yet been formally investigated.

To better characterise the sequential organization of actions during tool-use behaviours, and the extent to which such behaviours produce nonadjacent dependencies, we herein draw upon models of mutual information decay. Mutual information (MI) refers to the predictability of a given sequence element when the state of another element elsewhere in the sequence is known. MI often decreases between elements separated by greater distances; for example, looking out of a window at the weather one morning provides some information about the likely weather events that same afternoon. However, knowing the weather several days, or weeks prior would carry increasingly less information about the likely weather conditions in the upcoming afternoon. As non-hierarchical, ‘regular’ grammar systems (such as reflexive chains, and Markov models) produce sequences based on neighbouring probabilistic dependencies, they exhibit exponential decreases in MI between elements separated by increasing sequential distances (Lin and Tegmark, 2017). Alternatively, hierarchical grammar systems are able to readily produce long-distance dependencies between elements, and as such, show higher MI scores between distantly separated elements, which decay following a power-law dynamic (Sainburg et al., 2022, 2019). Hierarchical and non-hierarchical structuring mechanisms may also be combined to form a composite structure (Sainburg et al., 2022, 2019), where shorter, non-hierarchical strings are hierarchically organized to produce longer sequences. Sequences produced by composite structuring may then display MI decay dynamics that are best approximated by exponential decay for sequential elements in close proximity, and power-law decays for elements separated further across sequences. MI decay dynamics have been used to evaluate corpora of human and non-human vocal behaviours (Sainburg et al., 2022, 2019), as well as the sequential actions of humans, zebrafish and fruit flies (Sainburg et al., 2020). However, they have not thus far been applied to the sequential tool-use behaviours of a nonhuman animal.

In the present study, we employ MI decay models to investigate two overlapping research questions, concentrating on the sequential organization of serial action in chimpanzee tool use:

[1] Do chimpanzees produce nonadjacent dependencies within their sequential actions during tool-use behaviours?
[2] What do the dynamics of MI decay in chimpanzee tool-action sequences suggest about their underlying grammars of production (Markovian, hierarchical, or a composite system)?

To answer these questions, we focus on the sequential organization of actions used by wild West-African chimpanzees (*Pan troglodytes verus*) when engaging in one of their most complex natural tool-use behaviours: the cracking of hard-shelled nuts with hammer and anvil stones. Through MI estimations, we demonstrate that the sequences of actions performed during tool use by two thirds of all sampled adult chimpanzees possess sequential dependencies at significantly greater distances than those predicted by simulated Markov models. For half of adults, this relationship remained once repeating actions were accounted for, suggesting that repeating actions alone could not account for dependencies at long distances for many adults. We also report that the sequences of actions used by the majority of chimpanzees during nut cracking yield MI decay profiles that are concurrent with previous qualitative descriptions of hierarchical structuring in the tool action of apes. We also note that there is detectable inter-individual variability in the distances at which dependencies occur in action sequences, and the most likely mechanisms of sequence generation detected by MI decay. We therefore report that whilst hierarchical organization is important for many chimpanzees, it is unclear whether it is used universally by all individuals. Thus, our findings provide novel insights into non-human apes’ capacity to draw upon supraregular rules of sequence production, and produce nonadjacent dependencies during a natural behaviour: tool use. In extension, our findings feed into the wider discussion surrounding the hierarchical abilities of animals, as well as possible homologous origins of hierarchical action processing between humans and our sister species.

## 2. Results

### 2.1 Sampled Sequence Data

We analysed a total of 251.85 min of video data of chimpanzees engaging in nut cracking, taken from the 2011/2012 field season, yielding action sequences from nine individuals (see Table 1). Within our analyses, actions were compound descriptions of both the functional movement produced by an individual, and the focal object of the action (e.g., “Place Nut” = 1 action - see the methods section *behaviour coding* for further information). Of these nine individuals, one (Velu) had fewer than 200 actions coded throughout the entire series of sampled videos. As entropy estimation from small sample sizes can be subject to high levels of error (Grassberger, 2003), this individual was omitted from our MI analyses. Subsequently, eight individuals were included in our MI analyses: two adult females (Fanle & Jire); four adult males (Foaf, Jeje, Peley & Tua); one juvenile female (Joya) and one infant male (Flanle).

**Table 1.** Summary of sequence data collected from the 2011/2012 field season at Bossou (including all complete and incomplete sequences). Eight individuals were included in MI decay modelling. ‘Time Coded’ relates to the total duration of time individuals were seen to be engaging with nuts, nut fragments and stone tools throughout all video footage. An ‘Element’ is defined as each action coded within a sequence, whereas ‘Action Types’ represent the different types of coded actions in a sequence (e.g., “Grasp Nut” “Place Nut” “Grasp Nut” – Number of elements: 3, Number of Action Types: 2). ‘Nuts Cracked’ marks the number of nuts observed to have been cracked by an individual across all available video footage. Mean Sequence Length Per Nut refers to the mean number of elements in a sequence that is directed at cracking open each individual nut. SE for all means were calculated via the normal approximation of the Poisson distribution. Ages marked with * are estimates, as these individuals were already present at Bossou when long-term monitoring began in 1976.

| Individual | Sex | Age (years) | Sampled for MI Analysis | Time Coded (MM:SS) | Number of Sequences | Sequence Elements (Total) | Mean Sequence Length ± SE | Number of Action Types | Nuts Cracked | Mean Sequence Length Per Nut ± SE |
| --- | --- | --- | --- | --- | --- | --- | --- | --- | --- | --- |
| Fanle | F | 14 | ✓ | 44:55 | 21 | 1368 | 65.1 ± 1.8 | 34 | 85 | 13.9 ± 0.4 |
| Flanle | M | 4 | ✓ | 24:03 | 35 | 453 | 12.9 ± 0.6 | 59 | 1 | 48 |
| Foaf | M | 31 | ✓ | 07:22 | 8 | 299 | 37.4 ± 2.2 | 18 | 10 | 31.2 ± 1.8 |
| Jeje | M | 14 | ✓ | 40:33 | 18 | 1373 | 76.3 ± 2.1 | 36 | 39 | 29.4 ± 0.9 |
| Jire | F | 53* | ✓ | 19:27 | 12 | 482 | 40.2 ± 1.8 | 33 | 22 | 16.0 ± 0.9 |
| Joya | F | 7 | ✓ | 36:52 | 39 | 1178 | 30.2 ± 0.9 | 55 | 22 | 36 ± 1.3 |
| Peley | M | 13 | ✓ | 34:05 | 15 | 1168 | 77.9 ± 2.3 | 44 | 45 | 21.7 ± 0.7 |
| Tua | M | 54* | ✓ | 55:33 | 15 | 1940 | 129.3 ± 2.9 | 43 | 79 | 22.4 ± 0.5 |
| Velu | F | 52* | ✗ | 01:07 | 2 | 16 | 8 ± 2 | 12 | 0 | 0 |

Across the eight individuals in our dataset, we recorded action sequences for the cracking of 303 individual nuts, using 8,261 individual actions, belonging to 82 unique classes of action (see Fig. 1 for a list of the most common classes of actions used by chimpanzees, and example sequences of actions performed during nut cracking). Flanle, the infant male, used the greatest variety of different actions during nut cracking (59 unique classes of action), followed by Joya, the juvenile female (55 unique classes of action). Both of these individuals interspersed short sequences of solo object play into their attempts at nut cracking, e.g. mouthing the hammer stone, coded as either ‘kiss HAMMER’ or ‘bite HAMMER’, depending on the force applied. For both of these individuals, we retained these actions as part of our coded action sequences, to avoid subjective bias of when individuals were or were not playing.

**Figure 1.**
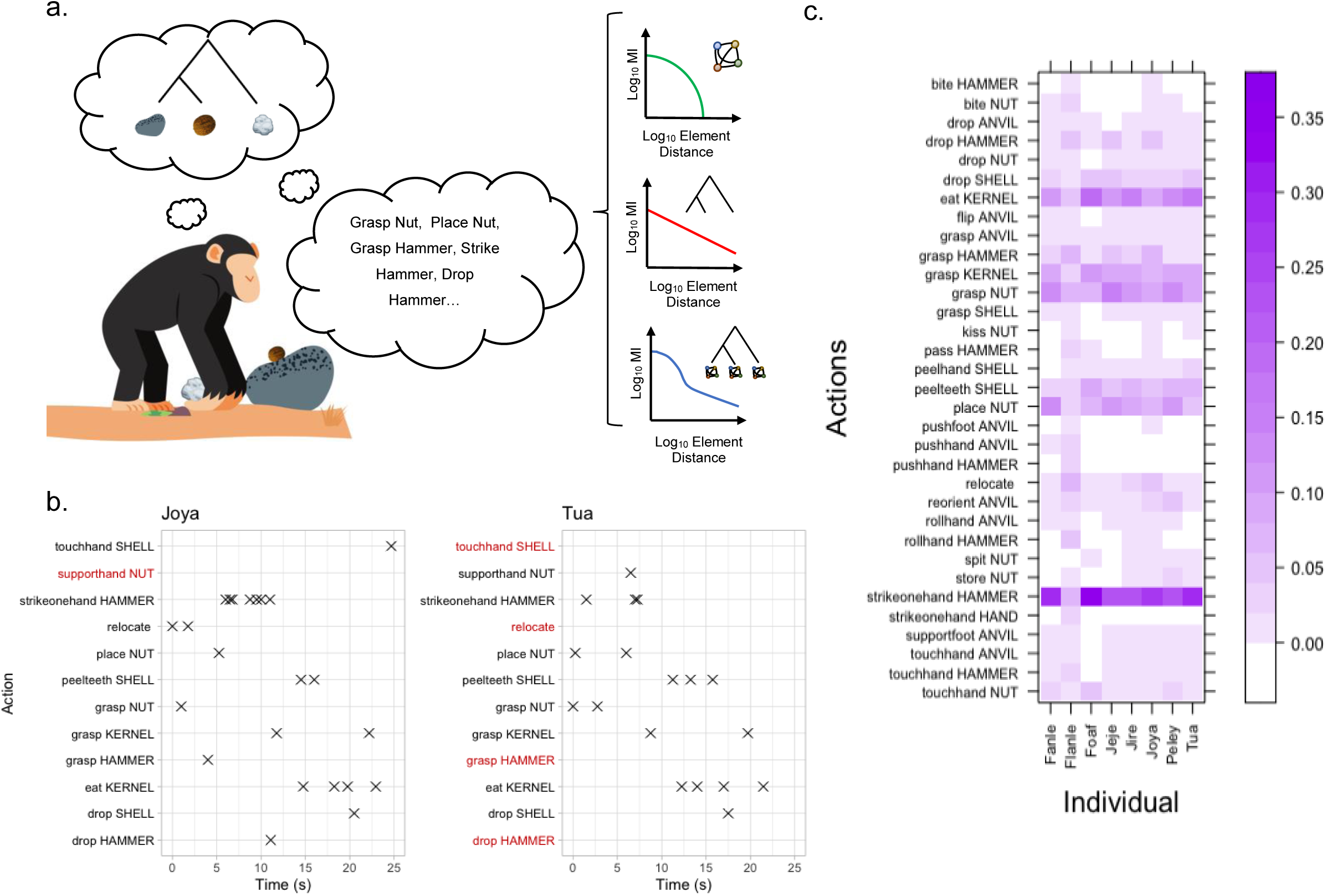
MI Decay models and the diversity of tool-use actions employed by chimpanzees during nut cracking. **(a)** A chimpanzee using two stone tools to crack open a nut (tree structure diagram represents hierarchical association of objects - see Matsuzawa 1996), and a complementary sequence of actions to facilitate object associations. To the right of the action sequence, three candidate structuring mechanisms are outlined, alongside their characteristic MI decay profiles (from top to bottom: purely Markovian, purely hierarchical, a composite of a hierarchical structure with Markov models at the terminal nodes). **(b)** Two representative behaviour time-plots showing the sequence of actions used by Joya (left) and Tua (right) to crack and consume a nut. Crosses mark where an action is coded as a discrete time point. Red action labels indicate where an action-type was not used by an individual when cracking and consuming each specific nut; however, they do not preclude these actions from being used by either individual when processing other nuts, and do not represent all possible actions. **(c)** A heatmap showing the proportion of different action types in the total corpus of action-sequence data for each individual (restricted to action types which constituted at least 1% of total observed actions for at least one individual). The darker purple shading means action types constitute a greater proportion of the corpus for each individual.

Adults used a smaller repertoire of actions when engaging in nut cracking. This repertoire included a small subset of high-frequency actions to facilitate nut cracking, with the five most common actions including ‘Strikeonehand Hammer’ (24.6%), ‘Eat Kernel’ (11.8%), ‘Grasp Nut’ (10.5%), ‘Place Nut’ (8.6%), and ‘Grasp Kernel’ (8%; in total, 63% of all actions within our dataset fell into one of these five action types). In addition to these frequently performed actions, an extended number of actions were performed less frequently, only where necessary, e.g. passing the hammer between the hands (pass HAMMER), or rotating the anvil stone to confer stability (reorient ANVIL; see Fig. 1c for more examples).

### 2.2. Maximum Inter-Element Distance MI Detected

To understand the extent to which chimpanzees produced non-adjacent dependencies in their sequential actions, we identified the maximum sequential distance at which dependencies occurred. For this analysis, we looked at the maximum sequential distance at which we detected positive MI values. We used an MI score which was adjusted for the occurrence of chance relationships between actions (MI_Adj_). For each individual, we compared this result to the maximum distances MI was positive in 100 sequences simulated from Markov models. This allowed us to determine whether dependencies in observed action sequences occurred at greater distances than those which can be produced through Markovian chaining of actions.

On average, the maximum sequential distances where MI_Adj_ remained above zero was 30.3 elements (N = 8, SD = 29.9; see table 3). For Flanle, the infant male, MI_Adj_ did not fall to zero before the maximum distance of 100 sequence elements. When excluding Flanle from this average, for whom the maximum distance over which dependencies were detected was over two standard deviations higher than all other individuals, the mean maximum sequential distance at which MI_Adj_ was positive was 20.3 elements (N = 7, SD = 10.8).

For six individuals (Fanle, Flanle, Jeje, Joya, Peley and Tua), a positive MI_Adj_ was detected at sequential distances that were significantly greater than those found in counterpart Markovian sequences (identified by a higher inter-element distance where MI_Adj_ was positive compared to the upper 95% CI of the Markov distribution; see Fig. 2 and Table S3). This result confirms that nonadjacent dependencies for these individuals occurred at distances greater than could be readily generated by comparative Markov models. Alternatively for two individuals, (Foaf & Jire), there was no difference between the maximum distance MI_Adj_ was detected in the observed action sequences compared to sequences simulated from Markov models.

**Figure 2.**
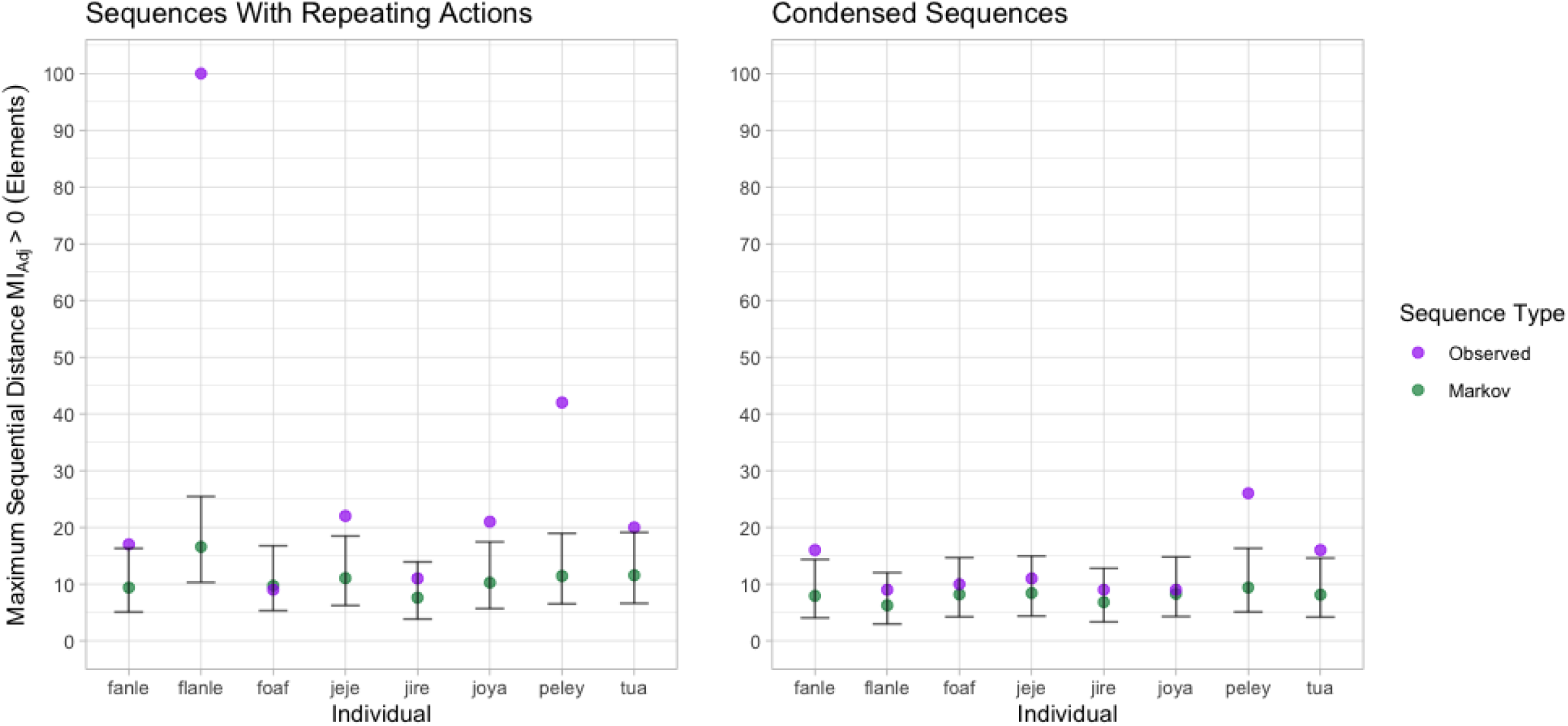
The maximum sequential distance dependencies were detected in action sequences and sequences from Markov models. Determined by the maximum distance that MIAdj was estimated to be greater than zero. MIAdj estimated from action sequences is in (purple), and estimates from sequences generated by an individual’s corresponding Markov models (green). The panel on the left is for data which includes repeating actions; the panel on the right corresponds to condensed action-sequence data. Bars represent a 95% confidence interval around estimated means for Markovian sequence data.

To determine whether successive repeats of elements (e.g. repeatedly striking a hammer) influenced the maximum inter-element distance where MI_Adj_ was above zero, observed action sequences were passed through a condensing algorithm that removed repeated elements, and MI_Adj_ was estimated in the same way as before. Separate Markov models were also constructed from condensed data, and our analysis was repeated identically to before.

For the condensed action sequences, the mean maximum sequential distances where MI_Adj_ was detected was 13.3 elements (N = 8, SD = 5.9). Flanle, the infant male, showed the largest decrease in the maximum distance where MI_Adj_ was greater than zero, reducing from possibly over 100 elements down to nine sequence elements. This implies that dependencies were heavily influenced by repeating actions in Flanle’s action sequences, possibly through repeated play actions.

For three individuals (Fanle, Peley and Tua), the maximum distance at which MI_Adi_ was positive remained significantly greater than sequences produced by their corresponding Markov models (see Fig. 2; see Table S3). These three individuals represent half of all adults in our sample. Following sequence compression, three individuals no longer exhibited dependencies at sequential distances greater than simulated Markov sequences: Flanle (infant male), Joya (juvenile female) and Jeje (adult male).

### 2.3 MI Decay Analysis: Observed Action Sequences

To understand how sequences of behaviours are generated, it can be informative to assess how mutual information decays between sequence elements which are progressively further apart (Lin and Tegmark, 2017; Sainburg et al., 2022, 2020, 2019). We looked at how MI_Adj_ decayed between actions which were increasingly separated in the action sequences produced by chimpanzees. Exponential, power-law, and composite decay models were all fitted to each decay profile (see Fig. 3), and the performance of each model in characterising MI_Adj_ decay was evaluated using AICc (where lower values represent a better explanation of the data, relative to the number of model parameters).

**Figure 3.**
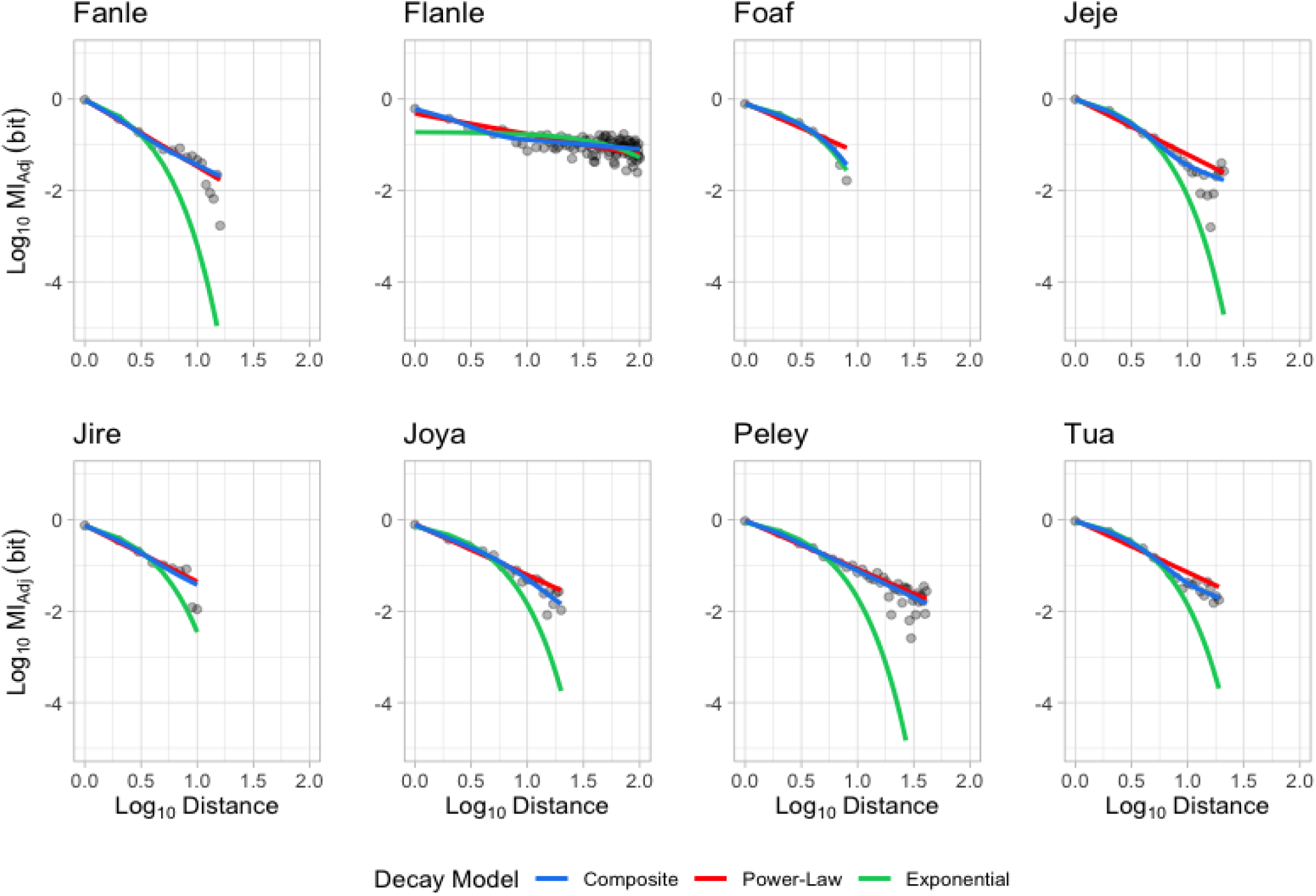
MIAdj decay profiles for each individual. The three candidate decay models are fitted. The exponential model is in green; the power-law model is in red, and the composite model is in blue.

For seven out of eight individuals, MI_Adj_ decay was best characterised by a model which included a power-law decay dynamic, indicating some aspect of hierarchical organization is involved in action-sequence generation (see Fig. 3 and Table 3; see Table S4 for AICc values). The MI_Adj_ decay profiles of two individuals (Fanle and Jire) were best explained by a purely power-law model (indicative of sequences produced by solely hierarchical structuring); while for five individuals (Flanle, Jeje, Joya, Peley and Tua), the optimal decay model was identified as the composite model, which combines periods of exponential and power-law decay. These composite models suggest that these five chimpanzees use a two-stage process of generating short subroutines through chaining together a handful of actions, before hierarchically organizing these subroutines into longer sequences. For one individual (Foaf), the exponential model was identified as the optimal model, implying that Foaf organized his actions using non-hierarchical means.

Similarly to our previous analysis of the maximum inter-element distances where we detected positive MI_Adj_, we repeated the analysis above on condensed sequences to ensure power-law relationships were not simply a product of highly repetitive elements (see Table 2). When analysing condensed action sequences, we were able to identify an optimal decay model for the action sequences of six individuals. Of the eight individuals in our analysis, the action sequences of five individuals were best explained by a model which included a period of power-law decay (two by a purely power-law, and three by a composite model), indicating partial or wholly hierarchical structuring. For two individuals (Jeje and Jire), multiple models offered competing explanations of decay dynamics: for action sequence data collected from Jeje, the composite and exponential models offered competing explanations of the data; whereas for Jire, the power-law and exponential models exhibited similar explanatory power. For these individuals, the extent to which hierarchy was involved during action-sequence generation was ambiguous. For the MI_Adj_ decay profile for Foaf, the exponential model continued to offer the best explanation of the data following sequence compression, implying non-hierarchical means of action-sequence generation.

**Table 2.** AICc scores for MIAdj decay models, derived from condensed sequences. For each individual, Max Element Distance highlights the greatest element distance included in the MIAdj datasets, and models are given in the order of preference (as determined by AICc – rows in bold indicate highest preferences). K represents the number of parameters estimated in each model. AICc describes the AIC score corrected for smaller sample size (see section 4.6). ΔAICc represents the difference between a given model’s AICc score and the optimal model’s AICc score for each individual. AICc.Wt describes the proportion of the total predictive power of all three models contained in each model, and Cum.Wt describes the accumulation of predictive power in order of model preference. LL represents the log-likelihood of each model given the data - AICc is estimated using both K and LL.

| Individual | Max Element Distance | Model | K | AICc | $\Delta AICc$ | AICc.Wt | Cum.Wt | LL |
| --- | --- | --- | --- | --- | --- | --- | --- | --- |
| Fanle | 15 | **Composite** | **5** | **-70.5** | **0.0** | **0.93** | **0.93** | **43.6** |
|  |  | Power-Law | 3 | -65.2 | 5.3 | 0.07 | 1.00 | 36.7 |
|  |  | Exponential | 3 | -47.2 | 23.3 | 0.00 | 1.00 | 27.7 |
| Flanle | 8 | **Power-Law** | **3** | **-18.5** | **0.0** | **0.98** | **0.98** | **15.3** |
|  |  | Exponential | 3 | -10.3 | 8.2 | 0.02 | 1.00 | 11.2 |
|  |  | Power-Law | 5 | 0.8 | 19.4 | 0.00 | 1.00 | 19.6 |
| Foaf | 9 | **Exponential** | **5** | **-36.7** | **0.0** | **1.00** | **1.00** | **23.8** |
|  |  | Power-Law | 3 | -22.7 | 14.1 | 0.00 | 1.00 | 16.8 |
|  |  | Composite | 3 | -14.1 | 22.6 | 0.00 | 1.00 | 22.1 |
| Jeje | 10 | **Composite** | **5** | **-43.0** | **0.0** | **0.72** | **0.72** | **34.0** |
|  |  | **Exponential** | **3** | **-41.2** | **1.8** | **0.28** | **1.00** | **25.6** |
|  |  | Power-Law | 3 | -23.0 | 20.0 | 0.00 | 0.00 | 16.5 |
| Jire | 8 | **Power-Law** | **3** | **-22.7** | **0.0** | **0.54** | **0.54** | **17.4** |
|  |  | **Exponential** | **3** | **-22.4** | **0.3** | **0.46** | **1.00** | **17.2** |
|  |  | Composite | 4 | 10.7 | 12.0 | 0.00 | 1.00 | 25.4 |
| Joya | 8 | **Power-Law** | **3** | **-26.5** | **0.0** | **1.00** | **1.00** | **19.2** |
|  |  | Exponential | 3 | -15.0 | 11.5 | 0.00 | 1.00 | 13.5 |
|  |  | Composite | 5 | -2.3 | 24.2 | 0.00 | 1.00 | 21.2 |
| Peley | 26 | **Composite** | **5** | **-121.8** | **0.0** | **0.80** | **0.80** | **67.5** |
|  |  | Power-Law | 3 | -119.1 | 2.7 | 0.20 | 1.00 | 63.1 |
|  |  | Exponential | 3 | -80.0 | 41.8 | 0.00 | 1.00 | 43.6 |
| Tua | 16 | **Composite** | **5** | **-92.2** | **0.0** | **1.00** | **1.00** | **54.5** |
|  |  | Exponential | 3 | -70.7 | 21.5 | 0.00 | 1.00 | 39.4 |
|  |  | Power-Law | 3 | -49.0 | 43.3 | 0.00 | 1.00 | 28.6 |

### 2.4 Transition Points for Composite Models

For each individual for whom our analysis showed a preference for the composite decay model, we also identified the transition point between primarily exponential and primarily power-law decay dynamics through the second differential of the curve in log-space (see Fig. 4.; see Methods section *Estimating Transition Points in Composite Model*). The maximum sequential distance prior to the transition between exponential and power-law decay dynamics ranged from 2-8 elements (mean = 4.6 elements, N = 5; see Table 3; see Fig. S4 for transition point profiles for each individual whose MI_Adj_ is best explained by the composite model). This implies that for individuals whose sequence structure is best approximated by composite models, non-hierarchical chaining of actions can be used to produce subroutines of 2-8 elements apart. These subroutines of actions are then hierarchically combined to produce extended behavioural sequences.

**Figure 4.**
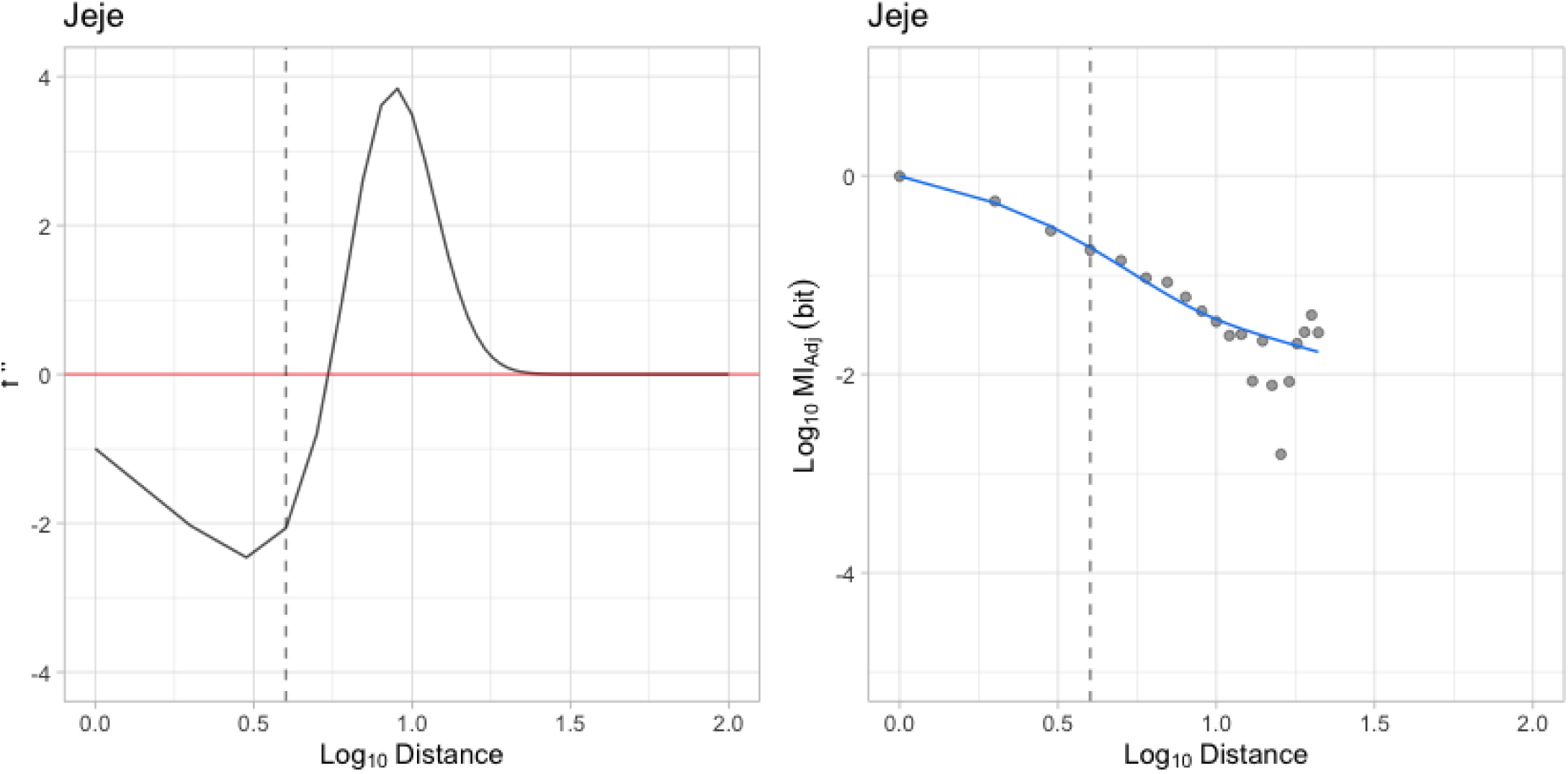
Example curvature analysis for the composite model. The panel on the right shows the composite model describing MI decay for Jeje. The panel on the left illustrates the second differential of this composite function. The point after the minimand of the second differential indicates the transition to power-law decay (indicated by a vertical dashed line). This point is mirrored on the plot on the right by the same line.

**Table 3.** Summary of maximum comparative distances, model preferences, and transition points for action sequences with repeating actions. Transition point describes the element distance where the composite model switches from a primarily exponential decay dynamic to one which is better generalised by a power-law. For each individual, the transition point, and mean number of actions used to crack open individual nuts, is provided. Total sample size for nut sequence length (i.e., number of nuts cracked throughout the study period) is included in brackets.

| Individual | Sex | Age Class | Maximum Sequential Distance $MI_{Adj} > 0$ | Preferred Model | Transition Point | Mean Sequence Length Per Nut ± SE (no. of nuts cracked) |
| --- | --- | --- | --- | --- | --- | --- |
| Fanle | F | Adult | 16 | Power-Law | - | 13.9 ± 0.4 |
| Flanle | M | Infant | 100 | Composite | 2 | 48 (1) |
| Foaf | M | Adult | 8 | Exponential | - | 31.2 ± 1.8 |
| Jeje | M | Adult | 21 | Composite | 4 | 29.4 ± 0.9 (39) |
| Jire | F | Adult | 10 | Power-Law | - | 16.0 ± 0.9 |
| Joya | F | Juvenile | 20 | Composite | 8 | 36 ± 1.3 (22) |
| Peley | M | Adult | 41 | Composite | 2 | 21.7 ± 0.7 (45) |
| Tua | M | Adult | 19 | Composite | 7 | 22.4 ± 0.5 (43) |

### 2.5 MI Decay Analysis: Markovian Sequences

To understand the rate at which our method would identify power-law decay dynamics in non-hierarchical sequences (akin to a false-positive rate for identifying signatures of hierarchies from non-hierarchical sequences), we simulated action sequences from Markov models, and ran the same model selection process as previously described. We created two ‘community Markov models’ (trained on action-sequences from all individuals) to act as a negative control. One was trained on all observed action-sequences, and another on the condensed version of all action-sequence data where repeating actions were removed. We generated 500 sequences from each of these models, estimated MI across the sequences, before fitting and evaluating the suitability of our three candidate models (exponential, power-law and composite). We estimated a false positive rate by identifying the proportion of sequences whose MI decay profiles were best described by models which contained power-law relationships.

For the Markov model trained on the entire corpus of action-sequence data, including repeating elements, all three decay models were successfully fitted for the MI_Adj_ decay profiles for 425 sequences. There was a significant, positive relationship between with the maximum sequential distance at which MI_Adj_ was detected, and the likelihood of successfully fitting all three models (Binomial GLM with Logit Link: Success ∼ Maximum Sequence Length, N = 425; Intercept: −9.25; Maximum Length: 1.53; Z = 8.96; p< 0.001), and this relationship also held for the condensed data (see table S5). Of the 425 model comparisons, AICc identified a single best model for characterising MI_Adj_ decay for 340 datasets (80% of all model comparisons). 52.6% of model comparisons identified the exponential model as highly competitive (see Fig. 5). In 37.6% of model comparisons the exponential model was identified as the sole best performing model, and in 15% of comparisons the exponential model was identified as one of multiple equally optimal models. 16.5% of comparisons showed a sole preference for the power-law model, and 25.9% of comparisons showed a sole preference for the composite model.

**Figure 5.**
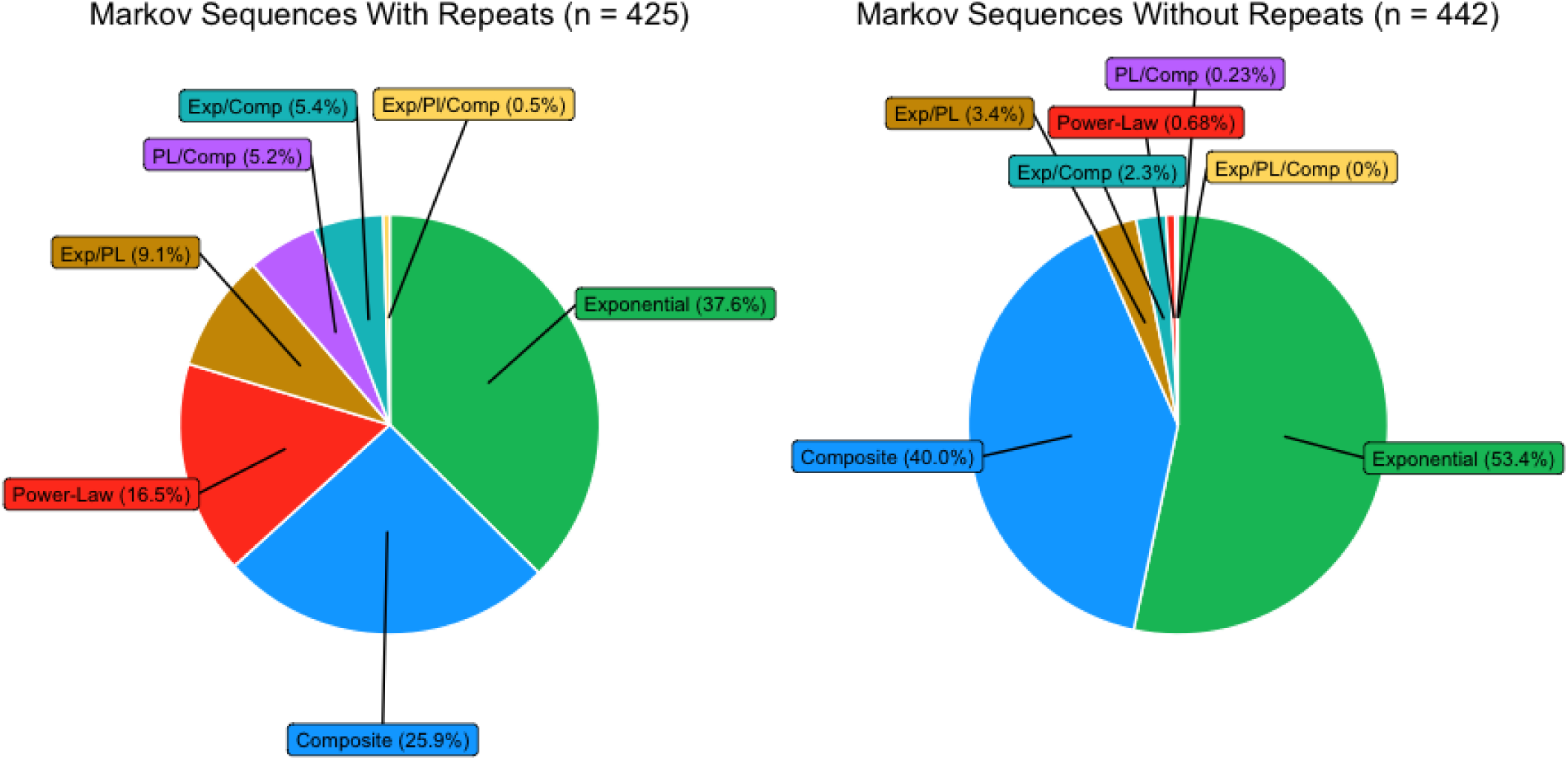
The proportion of instances MIAdj decay was preferred when generating sequences from community Markov models. The panel on the left relates to 425 sequences generated from a Markov model which can produce repeating actions in a sequence. The panel on the right relates to 442 sequences generated from a Markov model which cannot produce repeating actions.

This rate suggests that, if chimpanzees were organizing sequences similar to non-hierarchical Markov models, we would expect composite and power-law models to be the sole best performing model in 42.4% of instances (approximately 3 of 8 individuals). In the action sequence data, this value was over double (7 individuals). Our simulation also predicts that, if generated through non-hierarchical means, the exponential model would be the best performing model for 3 individuals. However, our analysis of the action-sequence data revealed that the exponential model was preferred for a single individual, Foaf.

A similar result was found when using a Markov model trained on condensed action sequences (see Fig. 5). If action sequences were generated through non-hierarchical means, we would expect composite and power-law models to be the sole best performing model in 40.68% of instances (3 of 8 individuals). We identified 5 individuals for whom composite and power-law models were the sole preferred model. Equally, we would expect the exponential model to be preferred in 53.4% of instances (4 individuals); however, much like the analysis for action-sequence data containing repeating actions, the exponential model was preferred for a single individual, Foaf.

## 3. Discussion

Hierarchical organization and nonadjacent dependencies are essential for the generation of many different elaborate sequential behaviours in humans (Chomsky, 2002; Dawkins, 1976; Fitch and Martins, 2014; Lashley, 1951; Mel’čuk, 1988; Rispens and Soto de Amesti, 2017; Rosenbaum et al., 2007; Wilson et al., 2020). However, these cognitively demanding aspects of sequential behavioural organization appear to be sporadically distributed across other animal taxa, and when present, may be restricted to specific behaviours. Given their close phylogenetic proximity to humans, characterizing how great-apes organize sequential behaviours is informative for understanding how human sequential behaviours evolved. This includes the evolution of cognition to support hierarchical organization of sequential behaviours, and the ability to trace and understand nonadjacent dependencies between elements in a behavioural sequence. The purpose of this study was to apply mutual information measures (estimates of dependency strength between sequence elements) to: (1) determine whether chimpanzees routinely produce nonadjacent dependencies between sequential actions during tool use, and if so (2) to determine whether the distance at which nonadjacent dependencies occurred in action sequences, and the dynamics of mutual information decay between increasingly separated elements, supports existing claims that chimpanzees hierarchically organize their action sequences during tool use. To answer both of these questions, we coded 8261 actions of eight wild chimpanzees at Bossou, Guinea, as they spontaneously cracked 303 individual nuts using stone tools.

For six out of eight of individuals, including two-thirds of adults (we take adulthood to be a proxy for experience and proficiency in tool use), nonadjacent dependencies were identifiable at sequential distances which significantly exceeded those produced in sequences simulated from Markov models. Thus, our data suggest that between the sequential actions produced by multiple chimpanzees during stone-tool use, nonadjacent dependencies occur at distances which exceed those found in comparable non-hierarchical sequences. For half of adults, these dependencies could not be solely explained by repeating actions; however for the other half of adults, as well as for both the juvenile and infant individual in our analysis, repeating actions contributed substantially to the emergence of nonadjacent dependencies. This finding does not preclude these individuals from generating some dependencies through hierarchical means (see further discussion on MI decay below); however, it does suggest that repeated actions contributed to a greater proportion of nonadjacent dependencies which spanned larger sequential distances. These results therefore suggest that there is detectable inter-individual variation in the means through which nonadjacent dependencies are generated within the action sequences of different chimpanzees, particularly between distantly separated actions.

Inter-individual differences in the distances at which dependencies occurred in action sequences is consistent with previously described differences in nut cracking techniques between chimpanzees at Bossou (Berdugo et al., 2023). Additionally, our analysis reveals that younger individuals may be more likely to produce dependencies through repeating actions, such as in the case of Flanle who showed a drastic decrease in the maximum distance dependencies detected after sequence compression. As the youngest individual in our analysis, Flanle was highly inexperienced in nut cracking, and we only observed Flanle successfully cracking open a single nut during the entire period of observation (for contrast, the other seven individuals for our analysis cracked a total of 302 nuts; mean = 43 nuts each, SD = 26.9). Flanle demonstrated the highest number of action types (59), indicating a greater diversity of different action types compared to the action sequences of adult chimpanzees during nut cracking (for whom action sequences were highly stereotyped in their use of a small number of different actions). This diversity of actions, as well as their repetition and flexible assembly, may have contributed to the anomalous distance at which dependencies were identified in the action sequences for Flanle. Further research is required to understand how these factors influence the emergence of sequential dependencies in animal behaviours.

Once repeating actions were controlled for, half of adult individuals in our analysis produced dependencies at distances which exceeded comparable Markov models. This result suggests that nonadjacent dependencies are produced by a substantial proportion of chimpanzees who are proficient in tool use, at distances which exceed expectations if sequences were generated through non-hierarchical means. Among the sequential behaviours of wild chimpanzees, tool use is the first domain in which such nonadjacent dependencies have been detected. This result provides new context to results from experimental paradigms which have previously revealed that captive chimpanzees are able to identify nonadjacent dependencies in both visual and auditory sequences (Sonnweber et al., 2015; Watson et al., 2020), by offering an ecologically-relevant domain of behaviour in which such cognition can be used. It is possible that, within the domain of manual actions, tool use is unique behaviour of wild chimpanzees which leads to the production of nonadjacent sequential dependencies. In another study performed on the sequential play behaviours performed by chimpanzees at Bossou, the accuracy with which an action in a play sequence could be predicted was aided by considering the two most recent actions, but actions further back in the sequence offered no additional benefit (Mielke and Carvalho, 2022). This suggests that play behaviours do not produce long-range dependencies, such as those we have identified during tool use. However, to validate whether tool use is unique in its production of nonadjacent dependencies, it will be necessary to analyse a wider range of sequential manual behaviours of wild chimpanzees. These analyses should include sequential manual behaviours which vary in their technical complexity, and should include both complex, multi-stage manual behaviours (e.g. nest building and complex foraging tasks which do not require tools), as well as simpler feeding tasks which require fewer processing steps (Gontier, 2024).

The sequential manual actions of non-human apes – including those used in tool use – have previously been described using qualitative frameworks which either explicitly or implicitly encompass hierarchical organization (Boesch et al., 2020; Byrne et al., 2013; Byrne and Russon, 1998; Carvalho et al., 2008; Estienne et al., 2017; Gontier, 2024); however, it has thus far been empirically challenging to exclude the possibility that these action sequences are produced by simpler structuring mechanisms (Vereijken and Whiting, 1998), particularly as human observers carry cognitive biases for identifying hierarchical structures in non-hierarchical sequences (Ferrigno et al., 2020; Fitch, 2014). Through characterizing MI decay across observed action sequences, we identified that for the majority of chimpanzees, MI decay included a period of power-law decay, indicative of hierarchical organization. Of this majority, the composite model was more frequently selected as the optimal model. This suggests that many chimpanzees were organizing actions into short subroutines through association rules between neighbouring elements, which were then hierarchically organized to produce longer sequences of tool-action. Through analysing the curvature of composite models, we identified that these subroutines of actions were between 2-8 elements long (mean = 4.6 sequence elements, N = 5). Our results are therefore consistent with previous descriptions of how great apes organize sequences of actions during food processing behaviours, including those which involve the use of tools (Byrne et al., 2013; Byrne and Russon, 1998; Gontier, 2024). Apes are theorized to decompose goals into subgoals which are addressed through short, stereotyped subroutines of actions (Byrne et al., 2013; Byrne and Russon, 1998), thus, these models of behavioural organization are identical to the composite structuring model in our analysis. Additionally, our analysis is able to objectively offer support to the hypothesis that apes organize sequences through the hierarchical organization of subroutines, without the presence of human bias for identification of hierarchical phenomena through subjective descriptions.

Despite the prevalence of the composite model in characterizing MI decay, our analyses did not provide conclusive results for all chimpanzees, as we could not resolve for a single best model for two individuals once we had accounted for repeating actions. For one individual, Foaf, the best model was identified to be the exponential model, which is associated with sequences which lack hierarchical organization. Such interindividual differences may be further evidence of differences in strategies for nut cracking behaviours between individuals, with a minority of individuals relying on alternative, non-hierarchical mechanisms of sequence generation. However, it is also important to recognize that for Foaf – the only individual for whom the exponential model was solely preferred – a much smaller corpus of actions was collected compared to those of other individuals (see Table 1). We are not able to say for certain whether sample size would have influenced the model preference for Foaf; however, given that sample size can influence entropy estimations (Grassberger, 2003), we can only conclude that this is a possibility for our analysis.

Whilst our analysis provides the strongest evidence for composite mechanisms of action organization in chimpanzee tool use, we deem it outside of the scope of this study to determine what parsing rules chimpanzees may use to facilitate a structuring system which relies on hierarchical and Markovian dynamics at different temporal scales (a limitation recognized in similar analyses on birdsong Sainburg et al., 2022, 2019). Additionally, our study is unable to provide insight into the exact representational mechanisms that chimpanzees use to produce successful sequences of tool action (Gruber et al., 2015). These limitations extend to both specific parsing procedures, and also the extent to which working memory is employed to account for distant dependencies. For example, it is possible that during nut cracking, physical traces of previous actions are maintained in the environment, rather than in the mind, reducing demands on memory (e.g. a nut positioned on top of an anvil would suggest it has already been placed there in a previous action; Byrne and Russon, 1998). However, our results do reaffirm that apes predominantly organize tool use behaviours using systems which encompass hierarchical organization. Further research is required to characterise the precise ‘grammars of actions’ used by chimpanzees to facilitate sequential behavioural organization during tool use, and for such questions, we anticipate that pattern-recognition algorithms will be particularly useful for identifying the precise subroutines of actions apes hierarchically combine to produce sequential behaviours (Gomez-Marin et al., 2016).

In contrast with other studies which employ analyses of MI decay (Sainburg et al., 2022, 2020, 2019), we used sequence data from Markov models to identify the rate at which power-law relationships would be identified from non-hierarchical sequences. This offered a false positive rate for misidentifying signatures of hierarchical organization. Power-law relationships were identified from non-hierarchical sequences at a non-negligible rate, indicating that MI decay analyses can produce false-positive results. However, this false positive rate was substantially lower than the rate at which power-law relationships were identified in observed action sequences. We therefore can conclude that signatures of hierarchical organization in chimpanzee action sequences did not represent false positives, but indicate systems of hierarchical sequential organization during action sequence generation. However, we strongly advise that future research employing MI decay analyses employ Markov models to estimate false positive rates of power-law detection. These false positive rates can then be compared with the rates of power-law detection from real-world sequential data to ensure that power-law model preferences are not statistical artefacts.

Using MI estimations, we present further evidence which affirms that the predominant structural system for chimpanzee tool use is one which encompasses hierarchical organization of subroutines. MI estimation has previously been used to detect hierarchical organization in the intricate grooming behaviours of Drosophila flies (*Drosophila melanogaster*) over an hour, and the swimming behaviours of Zebrafish (*Danio rerio*) during phototaxis paradigms (Sainburg et al., 2020), and for both species, the composite model was the best choice for explaining information decay over the action sequences of individuals. Our results suggest a possible homologous structuring system exists as a common method for chimpanzees to sequence actions in a highly complex natural tool-use behaviour: nut cracking. Additionally, the hierarchical complexity of early hominin tool-related action is believed to have increased throughout the course of hominin evolution, with the production of more ancient Oldowan tools requiring less complex hierarchical organization than the production of more recent Acheulian stone tools (Stout et al., 2021). Thus, the hierarchical organization of tool-use behaviours may have been an adaptation which was present in the evolutionary ancestor of humans and apes – stemming from more basal hierarchical systems of action – which was then elaborated across the course of human evolution into a state of higher complexity. Understanding the dynamics of this evolutionary transition would be greatly aided through further comparative research into the sequential organization of actions used by chimpanzees and humans during a greater variety of tool-use tasks, as well as everyday behaviours which require careful sequencing of manual actions (Gontier, 2024).

As previously mentioned, outside of tool use, the presence of hierarchical thinking in primates may also manifest within alternative behavioural contexts and social cognition, including in species other than chimpanzees (Bergman et al., 2003; Lameira et al., 2024; Mielke and Carvalho, 2022). In all of these instances, comparative research across domains of behaviour would be of interest, both within and between species. For example in apes, comparisons of chimpanzee play sequences and tool use may be informative for understanding when nonadjacent decencies can be expected in sequential manual actions, and how this expectation can be linked to fast-changing goals (such as in play behaviours) or goals which are sustained over extended periods of time (such as in tool use). Additionally orangutans offer an excellent opportunity to investigate whether the hierarchical complexity of vocal sequences, and the sequences of manual actions performed during tool use, covary between individuals. Orangutans would be particularly suitable for such an analysis, given their habitual use of hierarchically organized long calls (Lameira et al., 2024), and that they spontaneously manufacture and use tools in the wild (Van Schaik et al., 1996).

Further research comparing the dynamics of hierarchical behaviours within ape species will clarify the extent to which hierarchical cognition is domain general in species that are closely related to humans. This question is currently central to a number of hypotheses surrounding the evolution of cognition for hierarchical syntax and nonadjacent dependencies in human language. For example, the statistical scaffolding hypothesis (Sainburg et al., 2020) posits that long-range statistical dependencies in non-linguistic behaviours – and a generalised sensitivity to such behaviours – has led to a scaffold onto which language could evolve, where hierarchical syntax and semantics can be understood as later additions that exploit such structures and sensitivities. Similarly, the Cognitive Coupling hypothesis suggests that the pre-existing cognitive architecture responsible for hierarchical tool behaviours was coupled with cognitive machinery responsible for communication, where such coupling occurred in multiple steps across hominin evolutionary history, driven by the need to socially learn and transmit increasingly more complex tool behaviours (Kolodny and Edelman, 2018).

We believe that the findings of our study are consistent with, but not confirmatory of, both the statistical scaffolding hypothesis, and the cognitive coupling hypothesis. We have presented data herein that supports the presence of nonadjacent dependencies in the sequential action of chimpanzee tool use, as well as decay dynamics of MI that are best generalised by models associated with the hierarchical organization of action. Evidence for these structural features of sequential organization in chimpanzee tool use imply homologous origins with the sequential organization of actions in humans, suggesting that these structural features may also have been present in their last common ancestor. The emergence of highly hierarchical organization within communicative behaviours, and the resultant production of frequent nonadjacent dependencies in communication, may then have evolved as human-specific traits, following the evolutionary divergence of the human and *Pan* lineages. Further empirical investigation into the sequential organization of calls and gestures in chimpanzees and bonobos would offer further clarity on this matter.

In sum, we have demonstrated that MI decay models are useful for the study of sequential action organization in animal tool use. We have identified that adult chimpanzees produce nonadjacent dependencies within their sequential tool-use behaviours, and exhibit interesting inter-individual variability in the distances at which dependencies occur and in the means through which they are generated. These dependencies include those which cannot be explained by repeated actions, which were prominent in the action sequences used by half of all adult chimpanzees during nut cracking. Our results of MI decay also support previous descriptions that chimpanzees organize their actions during tool-use through the hierarchical organization of subroutines of actions. We therefore conclude that the tool-use behaviours produced by wild chimpanzees share many structural features with complex behaviours of humans, including for tool-use, language and music. Further research is required to understand the inter-individual variability we detected within our results. This includes the possibility that such structural features may not be universally employed by chimpanzees during tool use, as well as the possibility that the outlying individual in our analysis was an artefact of small sample size. Additionally, research comparing tool use with other hierarchical behaviours observed in primates will offer a useful lens through which to explore the domain generality of hierarchical cognition, as well as the possibility that hierarchical cognition was exapted for different behaviours across the evolutionary history of hominins.

## 4. Methods

### 4.1 Study Site

Bossou-Nimba (07° 390’ N; 008° 300’ W) is one of a small number of long-term field sites focused on the study of wild West-African chimpanzees (Matsuzawa et al., 2011). Located in South-Eastern Guinea, Bossou is surrounded by primary and secondary forest, and is home to a small community of wild chimpanzees whose behaviour and ecology has been studied systematically since 1976. The community’s home range covers an area of approx. 5-6 km (Sugiyama, 1981). Bossou chimpanzees are known to perform a wide variety of tool-use behaviours, including nut cracking (Biro et al., 2006, 2003; Carvalho et al., 2008; Hayashi, 2015; Inoue-Nakamura and Matsuzawa, 1997; Matsuzawa, 1994); leaf-folding (Tonooka, 2001), ant-dipping (Humle and Matsuzawa, 2002), and pestle-pounding (Yamakoshi and Sugiyama, 1995).

### 4.2 Data Collection

Data collection on nut-cracking behaviours is conducted at the “outdoor laboratory” at Bossou (Biro et al., 2006; Carvalho et al., 2008; Matsuzawa, 1994; see Fig. S1). Located in close proximity to the peak of Mount Gban, the outdoor laboratory is a natural rectangular clearing measuring approx. 7 × 20 m^2^ and operating as an in-situ experimental facility during the dry season of each year (November – February). Chimpanzees visit the clearing spontaneously as part of their daily ranging (at a rate that varies from once or twice a day on consecutive days, to returning after several days), and are provided with a selection of numbered stones of varying sizes, shapes and raw-material types. Chimpanzees are also provided with several piles of nuts, including naturally occurring oil-palm nuts (*Elaeis guineensis*), as well as Coula nuts, (*Coula edulis*), which despite not occurring naturally at Bossou, have been provided to chimpanzees at the outdoor lab since 1993 (Biro et al., 2006, 2003). A branch of oil-palm fruits is also provided, which is replenished frequently. Additionally, the outdoor laboratory features a tree trunk with a wide cavity inside, which acts as a water point. The outdoor laboratory is surrounded by thick vegetation on three sides, and on the fourth side an artificial wall made of palm fronds is used to allow human observers to observe chimpanzee behaviours out of sight, whilst simultaneously recording behaviours using tripod-mounted cameras. These conditions allow chimpanzee behaviours to be recorded in their natural habitat from close range but with minimal interference from human observers.

The several decades of video footage collected at Bossou have recently been compiled into a longitudinal video archive, spanning 1988 to the present (Schofield et al., 2019). As such, the Bossou video archive presents a unique opportunity to study wild chimpanzee behaviour in situ through both longitudinal and cross-sectional lenses. This study takes a cross-section of video footage data from the Bossou video archive, focusing on nut-cracking behaviours collected at the outdoor laboratory in the 2011/2012 field season.

### 4.3 Subjects

Since the start of systematic research at Bossou, the site is known to have supported a community of around 18-22 individuals; however, the population has been in decline since a disease outbreak in 2003 (Matsuzawa et al., 2004), and as of April 2022, they number only 6 individuals. In the 2011/2012 field season, the community consisted of 10 adult individuals (4 male and 6 female), 1 juvenile female and 2 infants males, with a community age range estimated to be from ∼ 1 to ∼ 54 years old (see Table S1). All available videos from the 2011/2012 field season were used during behaviour coding (see below), and all individuals who visibly interacted with a stone-tool, nut, or nut-fragment (including shells and kernels from their own or another’s previous nut-cracking event) whilst present in the outdoor laboratory were included in behaviour coding.

### 4.4 Behaviour Coding

Behaviour coding was conducted using BORIS (Behavioural Observation Research Interactive Software; Friard and Gamba, 2016). Digitised video recordings were passed to BORIS and visualised on a 59 cm monitor. An ethogram of codable behaviours was developed based on Inoue-Nakamura & Matsuzawa’s (1997) nut cracking ethogram (Inoue-Nakamura and Matsuzawa, 1997). Our ethogram included 34 codable actions, e.g. “Grasp”, “Place”, “Strike”, “Peel” etc. (see Table S2 for full ethogram) where each action was coded alongside one of a number of available objects, including: 1. Nut (uncracked), 2. Hammer, 3. Anvil, (Anvils were distinguished from Hammers on a case-by-case basis, based on how chimpanzees used each stone-tool), 4. Kernel – taken to be the edible inner part of the nut, 5. Shell – taken to be the inedible outer parts of the nut, 6. Bare Hand. In our analysis, each of these bigram descriptions (e.g. ‘Grasp Nut’) were considered to be an individual action. Some object-directed actions are continuous over time, and may occur simultaneously alongside other behaviours, e.g., a chimpanzee may support an anvil with its foot for an extended duration of time, whilst successively striking a nut with a hammer stone. To allow for analysis by mutual information decay, simultaneous actions were coded at the first point at which they were elicited; thus, all actions were coded as discrete-time points. This allows for actions to be described in one linear sequence (permitting MI decay analyses), whilst also retaining information surrounding the order in which chimpanzees began to externalize each object-directed action. Following action coding, each individual action was taken to constitute an individual sequence ‘elements’ in our analysis (i.e. “Grasp Nut” = 1 sequence element).

Action coding commenced when an individual began interacting with nuts, nut fragments or stone tools. Coding ceased when interacting with nuts and stone-tools to engage in another behaviour. Individuals who ceased interacting with nuts, nut-fragments and stone-tools for longer than one minute, but were not clearly engaging in another behaviour, were deemed to have begun resting, and behaviour coding stopped. Behaviour coding also ceased if an individual engaging in nut-cracking behaviours could no longer be clearly observed from the video footage, such as through manoeuvring their body to block their behaviours from being visible to cameras, moving out of shot of the cameras, or cameras panning away from focal individuals. Under instances where chimpanzees moved back into view following termination of a sequence due to poor visibility, behaviour coding resumed. Thus, behaviour coding produced a mixture of complete sequences, (where individuals were observed engaging in tool use from start to end) as well as shorter fragments of the tool use sequence. These shorter fragments contained the actions used to crack a subset of nuts within a longer sequence of cracking nuts, as well as shorter sequences of actions for raw material acquisition, and the disposal of tools and inedible shell.

Prior to MI analyses, sequences were concatenated in the order they were collected. Additionally, for any action sequences which included successful cracking of nuts, we counted the number of actions used by an individual to crack open a nut, consume all corresponding kernel, and dispose of any waste shell. We then estimated the mean number of actions performed to crack individual nuts for each individual, for comparison with transition points (see *Estimating Transition Points in Composite Models*).

Behaviour coding was conducted exclusively by EHS. To ensure our ethogram produced repeatable results, we ran interobserver reliability tests, which returned scores of 94-96% (see supplementary materials for further details).

### 4.5 Mutual Information Estimation

To estimate MI values at various element distances, sequences for each individual were concatenated in order of collection. Mutual information can be determined by the reduction in entropy of a point in a sequence, when the state of another sequence element is known. As such, information can be estimated through independent measures of entropy at different points in a sequence (the marginal entropy of points X and Y), and also the joint entropy of actions co-occurring together at a given sequential distance. To estimate information in our action sequences, marginal and joint entropies were estimated across concatenated sequences at distances ranging from 0 to 100 elements apart. MI was estimated at each distance as:

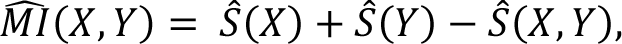

where X is the distribution of the initial elements; Y is distribution of elements n-items further along the sequence; 𝑆̂(𝑋) and 𝑆̂(𝑌) are the marginal entropies of X and Y respectively, 𝑆̂(𝑋, 𝑌) is the joint entropy of elements n distance apart, and 𝑀𝐼̂(𝑋, 𝑌) is the mutual information estimate for elements n distance apart. During entropy estimation, we followed the recommendations of Sainburg et al. (Sainburg et al., 2022, 2020, 2019) and Lin & Tegmark (Lin and Tegmark, 2017) by using the Grassberger method (Grassberger, 2003), which accounts for under-estimation of true entropy from finite samples:

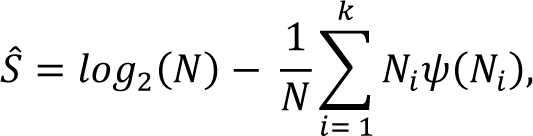

where 𝑆̂ is the marginal or joint entropy; N is the total number of elements in the distribution; K is the number of different groups of elements in the distribution, and 𝜓 is the digamma function. To account for lower bounds of estimated MI, sequences underwent 1000 pseudorandom permutations to generate randomised joint-entropy distributions. MI was then estimated using the mean of the permuted joint-entropy distribution, and the marginal entropy values from the in-sequence data, using the same equation as before. This MI estimate is taken to approximate MI measures which arise from our data due to chance relationships (Sainburg et al., 2022, 2020). The chance MI measure - 𝑀𝐼̂_𝑠ℎ_ (𝑋, 𝑌) - was subtracted from our data, to give an adjusted MI score which more accurately reflects the intentional structure of observed action sequences

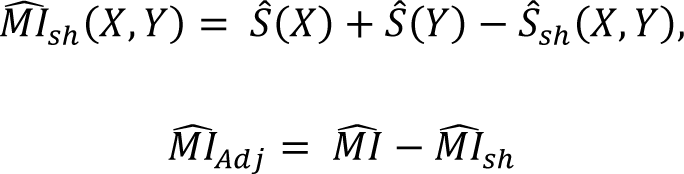

During MI estimation, it is possible for MI to decay to zero before reaching the maximal comparative distance of 100 elements. Therefore, we used the 1000 sequence permutations to construct 95% confidence intervals surrounding 𝑀𝐼̂_𝑠ℎ_ (𝑋, 𝑌) at each sequential distance between 0-100 elements. At each sequential distance between 1-100 elements, we took the upper value of the 95% confidence limit, and subtracted this value from the 𝑀𝐼̂ estimated from the original action-sequences at each corresponding distance:

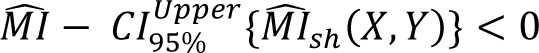

We took the first comparative distance this value fell below zero to indicate that estimated MI was no longer significantly different to what could be expected from chance relationships, and therefore could no longer be treated as different to zero. For MI decay profiles which decayed to zero before reaching the maximum comparative distance of 100 elements, MI decay data was restricted to include solely the comparative distances that preceded the complete decay of MI. For all subsequent analyses and modelling, the adjusted MI score was used (MI_Adj_).

### 4.6 MI Decay Model Fitting and Selection

Three models were used to characterise decay of adjusted MI scores: (1) an exponential model to approximate MI_Adj_ decay dynamics where sequences are produced by Markovian adjacent-dependencies (Lin and Tegmark, 2017; Sainburg et al., 2022, 2020, 2019); (2) a power-law model to approximate MI_Adj_ decay predicted by fully-nested hierarchical sequences (Sainburg et al., 2022, 2020, 2019); and (3) a composite model to approximate MI_Adj_ decay from sequences which utilise nested hierarchical structuring at higher orders of abstraction, before ordering elements through Markovian dynamics at local scales (Sainburg et al., 2022, 2020, 2019):

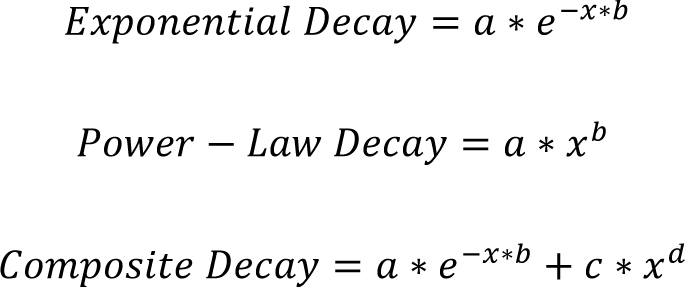

where x represents inter-element distance. As reducible and periodic Markov chains may produce MI_Adj_ scores which decay to positive constants (Lin and Tegmark, 2017), best-estimate Markov chains were modelled from sequence data for each individual, and checked to determine whether they were irreducible and aperiodic. Markov models were estimated in R using the *markovchainFit()* function from the *markovchain* package (Spedicato, 2017). Irreducibility and periodicity of these Markov models were then determined using the functions *is.irreducible()* and *period()* respectively. As all best-estimate Markov chains were identified as irreducible and aperiodic, we fitted our exponential model in the absence of a constant, allowing models of MI_Adj_ to decay towards zero. Both the power-law and composite decay models were also fitted in the absence of constants to also allow these models of MI_Adj_ to fall towards zero with increasing sequential distances (see the Quantification and Statistical Analysis section for more information on model fitting).

The fit of the three models were compared using an adaptation of the Akaike Information Criterion (AIC). AIC provides a numerical means to compare goodness-of-fit between models constructed on identical data-sets, whilst also penalising extra parameters used to construct the model. Such penalization of additional model parameters thus helps to reduce the likelihood of overfitting more complex models to their underlying data. Models with lower AIC scores are taken to give more accurate approximations of data, relative to the number of parameters used to fit them. To mirror similar analyses of MI decay on birdsong, language, and human action (Sainburg et al., 2022, 2020, 2019), we opted to compare model fits using AICc. AICc scores models similarly to AIC; however, AICc introduces an additional penalty for each model parameter, to correct for overfitting on smaller datasets (Cavanaugh and Neath, 2019; Sainburg et al., 2022, 2020, 2019)

As repeating elements in a sequence (e.g. successively striking a hammer) can produce nonadjacent sequential dependencies, we repeated the analyses outlined above – specifically in the sections *Mutual Information Estimation* and *MI Decay Model Fitting and Selection* – on a corpus of condensed sequences for each individual. Condensed sequences were produced by passing the concatenated sequence corpus for each individual through an algorithm which removed repeated actions are replaced them with a single codon (e.g. ‘Strike Hammer’, ‘Strike Hammer’, ‘Strike Hammer’, is replaced by a single codon: ‘Strike Hammer). By analysing this condensed corpus of sequences in parallel with observed sequences of chimpanzee tool action, we are able to separate out dependencies produced by repeated actions, and those produced by alternative means, including the hypothesised use of nested subroutines.

### 4.7 Estimating Transition Points in Composite Models

For those individuals whose MI_Adj_ decay profiles were best approximated by a composite model, we identified the point at which the model transitioned from exponential decay to power-law decay using the change in gradient of the curve in log-space (log_10_ MI_Adj_ and log_10_ element distance). Through finding the second differential of the curve in log-space, we took the minimand of this function to be the point immediately prior to where the function begins to switch to a power-law decay. The point at which the composite model switches to primarily a power-law decay can be considered to be the furthest inter-element-distance over which Markovian dynamics are likely to influence the structure of observed action sequences.

### 4.8 Markov Models and Simulated Sequences

Sequences produced from some Markovian grammars may produce dependencies which span over multiple elements (including short-range nonadjacent elements), especially when Markov models have highly predictable element transitions, leading to multiple elements being produced in highly stereotyped strings. It is therefore necessary to determine whether the distances over which sequential dependencies occur in chimpanzee tool-action are significantly greater to those that can be readily generated by equivalent Markov models. Using the Markov models generated from observed action sequence data from individual chimpanzees (see section above: *MI Decay Model Fitting and Selection*), we generated 100 Markov sequences matching the total corpus length for each individual. For all Markov sequences, we estimated MI_Adj_ Decay using the same method as outlined in the sections *Mutual Information Estimation* and *MI Decay Model Fitting and Selection*. From the 100 Markov Sequences for each individual, we constructed a probability distribution for the maximum sequential distances where dependencies could be detected, using the maximum sequential distance MI_Adj_ > 0 for each sequence. From these probability distributions, 95% confidence intervals were constructed using the exact method for Poisson distributions.

To determine whether dependencies occurred at greater distances in observed action sequences compared to those expected from Markovian structuring, we compared the maximum sequential distances where MI_Adj_ > 0 in observed action sequences versus to their corresponding 95% confidence interval for Markov sequences. This process was repeated for each individual using condensed sequence data, to determine whether the maximum distances over which dependencies were detected was influenced by repeating actions in both observed and Markov sequences.

To determine the extent to which Markovian approximations of chimpanzee tool-action produce MI_Adj_ decay profiles which are best approximated by exponential decay dynamics, we used two ‘community’ Markov models to produce control sequences. We concatenated sequence data from all eight individuals, and used this extended corpus to train an initial ‘community’ Markov model which reflected possible sequential-action combinations seen by all individuals within our analysis (similar to E-Languages in linguistics, which reflect the corpus of externalized utterances within a population; Chomsky, 1986). From this community Markov model, we simulated 500 sequences, each 1000 elements long, which were used for MI_Adj_ Decay analyses and Model fitting as outlined in sections *Mutual Information Estimation* and *MI Decay Model Fitting and Selection*. The proportion of MI decay profiles which were best characterized by exponential decay dynamics were calculated using AICc scores. This process was repeated with condensed sequence data, to produce a second community Markov model which approximated sequential relationships in the absence of repeating actions (see Fig. S5 for a graphical illustration of both Markov models). For MI Decay analyses from Markovian sequences, the University of Oxford’s Advanced Research Computing Facility was used to run MI Decay analyses in parallel (Richards, 2015).

### 4.9 Statistical Analysis

All statistical analyses were run in R Studio v.1.2.5001, using R v.4.2.1 (2022-06-23) – “Funny-Looking Kid” (R Core Team, 2022). For both mean sequence length (including sequences aimed at individual nuts, as well as sequences of continuous coding during data collection), and for the mean maximum sequential distances where MI_Adj_ > 0, 95% confidence intervals were estimated using the exact method for data following a Poisson distribution. Where standard errors are presented for these means, they were produced using the normal approximation of the Poisson Distribution. During MI_Adj_ estimation, permuted sequences were generated to estimate positive MI values produced by chance relationships, which could then be subtracted from MI values estimated from observed action sequences. MI at each sequential distance (0-100 elements) was averaged across all 1000 permuted sequences by finding the arithmetic mean. 95% confidence intervals about this mean were calculated using the standard error of MI at each sequential distance, and the following formula:

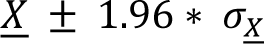

where 𝑋 corresponds to the arithmetic mean value at each sequential distance, and 𝜎_𝑋_ is the corresponding standard error.

All information surrounding estimation of transition matrices for Markov models can be found in the Method Details. The candidate MI decay models (exponential, power-law and composite) were fit to MI_Adj_ data using the nlsLM() function of the minpack.lm package (Elzhov et al., 2016), which uses the Levenberg-Marquardt algorithm to fit non-linear curves by least-squares. Following model fitting, MI estimates, predicted MI values, and inter-element distances were logged to base 10 to allow for decay dynamics to be visualised in log-space. AICc scores for each model were calculated in R using the Multi-Model Inference package (MuMIn; Bartoń, 2009).

## Author Contributions

E.H.S., D.B. and T.G. conceived and designed this project. D.B., M.H., and T.M. conducted and oversaw video data collection at Bossou. D.S. processed, cleaned and curated the archive. E.H.S. performed behaviour coding from video data, data analysis, and produced all figures. E.H.S., D.B. and T.G. discussed results and provided comments on the analysis. E.H.S. led the writing of this paper, with all authors contributing throughout drafting and revisions.

## Acknowledgments

E.H.S. thanks National Environmental Research Council (NERC) and United Kingdom Research and Innovation (UKRI) for financial support (grant ref. NE/L002612/1). T.G. thanks the Swiss National Science Foundation (SNSF) for financial support (grant ref. PCEFP1_186832). T.M. was supported by grants from Ministry of Education, Culture, Sports, Science and Technology, Japan (MEXT - #12002009, #16002001, #20002001, #24000001, #16H06283) and Japan Society for the Promotion of Science (JSPS) (Core-to-core CCSN and U04-PWS). M.H. was supported by grants MEXT/JSPS 15K00204, 19K21824, and JP17H06381 (Evolinguistics). D.S. was supported by grants from The Clarendon Fund, Boise Trust Fund, and Wolfson College, University of Oxford. We thank Susana Carvalho (University of Oxford, UK) for their help in curating the Bossou video archive, and Tim Sainburg (University of Harvard, USA) for statistical advice. We thank Anna Ahlberg and Melissa Birtles for offering their time to test the repeatability of our ethogram. The authors would like to acknowledge the use of the University of Oxford Advanced Research Computing (ARC) facility in carrying out this work. (http://dx.doi.org/10.5281/zenodo.22558). We are also grateful to all the KUPRI researchers who have helped collect data at Bossou in the past. Special thanks are due to Direction General de la Recherche Scientifique et l’innovation Technologique (DGERSIT) and the Institut de Recherche Environnementale de Bossou (IREB), République de Guinée, for assistance and research permission to conduct field work at Bossou; as well as research assistants Boniface Zogbila, Gouanou Zogbila, Henry Gbelegbe, Marcel Doré, and Gilles Doré for their help in the field.

