## Supplementary Materials for "Nonadjacent Dependencies and Syntactic Structure of Chimpanzee Action During a Natural Tool-Use Task"

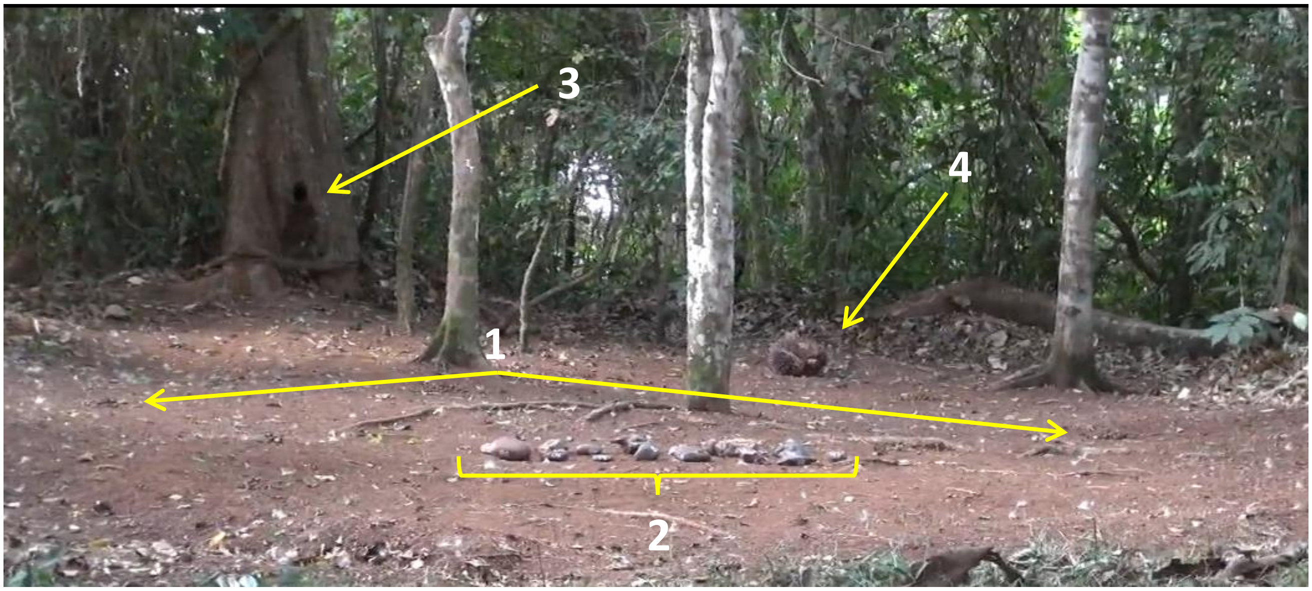


**Figure S1. The outdoor laboratory, Bossou**. Key features of the outdoor laboratory are: 1. Provided piles of oil-palm nuts (two piles are indicated; typically, a total of 6-12 piles of 40-50 nuts each were provided), 2. Central matrix of available stone tools, 3. Water point, 4. Bunch of oil-palm fruits.

**Table S1. Chimpanzees present at Bossou in the 2011/2012 field season.** Ages are listed as of Jan 2012.

| Individual | Sex | Age Category | Birth | Age | Relation | Hand use*** |
|---|---|---|---|---|---|---|
| Tua | M | Adult | 1957* | 54* | None | L |
| Foaf | M | Adult | 1980 late | 31 | Son of Fana | R |
| Jeje | M | Adult | 1997 Dec | 14 | Son of Jire | L |
| Peley | M | Adult | 1998 Apr | 13 | Son of Pama | L |
| Fana | F | Adult | 1956* | 55* | Mother of Fanle** | R |
| Jire | F | Adult | 1958* | 53* | Mother of Jeje | L |
| Velu | F | Adult | 1959* | 52* | None | R |
| Yo | F | Adult | 1961* | 50* | None | L |
| Pama | F | Adult | 1967* | 44* | Mother of Peley | — |
| Fanle | F | Adult | 1997 Oct | 14 | Daughter of Fana | R |
| Joya | F | Juvenile | 2004 Sep 2 | 7 | Daughter of Jire | L |
| Flanle | M | Infant | 2007 Sept 14 | 4y 4m | Son of Fanle | (L) in Dec |
| Fanwa | M | Infant | 2011 Nov 4 | 0y2m | Son of Fanle | — |

* Estimated age; these individuals were already present at Bossou when long-term study started in 1976.

** Fana is also the grandmother of Flanle and Fanwa; she took care of Flanle from the age of 4, following the birth of Fanwa.

*** Hand use refers to the hand (L, left or R, right) the individual uses to hold the hammer stone during nut-cracking. "—" refers to individuals that have never or not yet been seen to crack nuts using tools.

**Table S2. Ethogram of codable actions for observations of nut-cracking behaviours.** Actions in bold are coded alongside a corresponding object. There are 6 possible corresponding objects: 1. Nut, 2. Hammer, 3. Anvil, 4. Kernel, 5. Shell, 6. Bare Hand. Event codes in italics are used to denote the start and end of observable sequences. Coding commenced when individuals began interacting with stones, nuts, or nut-fragments. Coding ceased when individuals began engaging in a new behaviour e.g. play, grooming, eating oil-palm fruits. On the occasions where an individual moved out of clear sight of the video recording, or actions became obscured by an individual's body position, sequences were terminated with a 'Not Visible' codon, and marked as incomplete.

| Action | Description |
|---|---|
| **Bite** | Place object in mouth and bite with teeth. Differs from 'Eat' as there is no consumption. Differs from 'Store' as object moves in and out of mouth whilst chimp is stationary. Differs from 'Peel Teeth' as bite applies general force, whereas peel with mouth is to remove shell fragments from kernel dexterously. Differs from 'Kiss' as object enters inside the mouth. |
| **Brush** | Brush objects off the anvil. |
| **Drop** | Place object(s) on ground. |
| **Eat** | Consume object. Differs from bite and store as it requires successful ingestion. |
| **Flip** | Flip object over. |
| **Grasp** | Grasp an object and move off the ground. |
| **Kick** | Apply rapid, hard force on object with the foot, so that the object is displaced. |
| **Kiss** | Place object to lips or nose, but not inside of the mouth. |
| **Pass** | Pass object between hands. |
| **Peel Hand** | Peel shell off of kernel with hands. |
| **Peel Teeth** | Peel shell off of kernel with teeth. |
| **Place** | Place an object on an anvil. |
| **Provide** | Directly hand an object to another individual. |
| **Pull Foot** | Pull an object across the ground with foot. |
| **Pull Hand** | Pull an object across the ground with hand. |
| **Push Foot** | Push an object across the ground with foot. |
| **Push Hand** | Push an object across the ground with hand. |
| **Rake Foot** | Pull many objects towards oneself using foot/leg. |
| **Rake Hand** | Pull many objects towards oneself using hand/arm. |
| **Reorient** | Rotate object horizontally. |
| **Roll Foot** | Roll object along the floor with foot. |
| **Roll Hand** | Roll object along the floor with hand. |
| **Spit** | Let object fall from mouth or lips. |
| **Stomp** | Whilst standing, apply strong force to object with foot. |
| **Store** | Place object(s) in mouth for transportation. Separated from bite, eat, peel teeth as it is followed by transportation across the outdoor laboratory, before then being removed from the mouth intact. |
| **Strike One Hand** | Strike with one hand, (and associated object). |
| **Strike Two Hand** | Strike with two hands, (and associated object). |
| **Support Foot** | Support object with foot. |
| **Support Hand** | Support object with hand. |
| **Take** | Receive an object directly from another individual. |
| **Throw** | Throw object away from self, horizontally or vertically. |
| **Touch Foot** | Touch or grasp object without moving it using foot. |
| **Touch Hand** | Touch or grasp object without moving it using hand. |
| Relocate | Stand up and move to a new area; is immediately followed by another coding action. |
| Start | Start of a sequence. |
| End Bout | End of a sequence. |
| Not Visible | Individual moves out of view, and coding cannot continue. Differs from 'End Bout', as there is no evidence the individual has stopped engaging with stones, nuts or nut-fragments. |

### Interobserver reliability

We ensured that our ethogram could be used to generate repeatable action sequences through running interobserver reliability against two alternative observers who received two short training sessions with the ethogram. All observers (EHS and two volunteers) coded a five-minute clip containing three chimpanzees, and coded all actions performed by each individual. Actions were randomly sampled from the corpus generated by EHS (n = 50). We assessed if Observers 1 & 2 coded the same action within a window of ± 4 s. Observer 1 matched on 96% of actions, and observer 2 on 94% of actions.

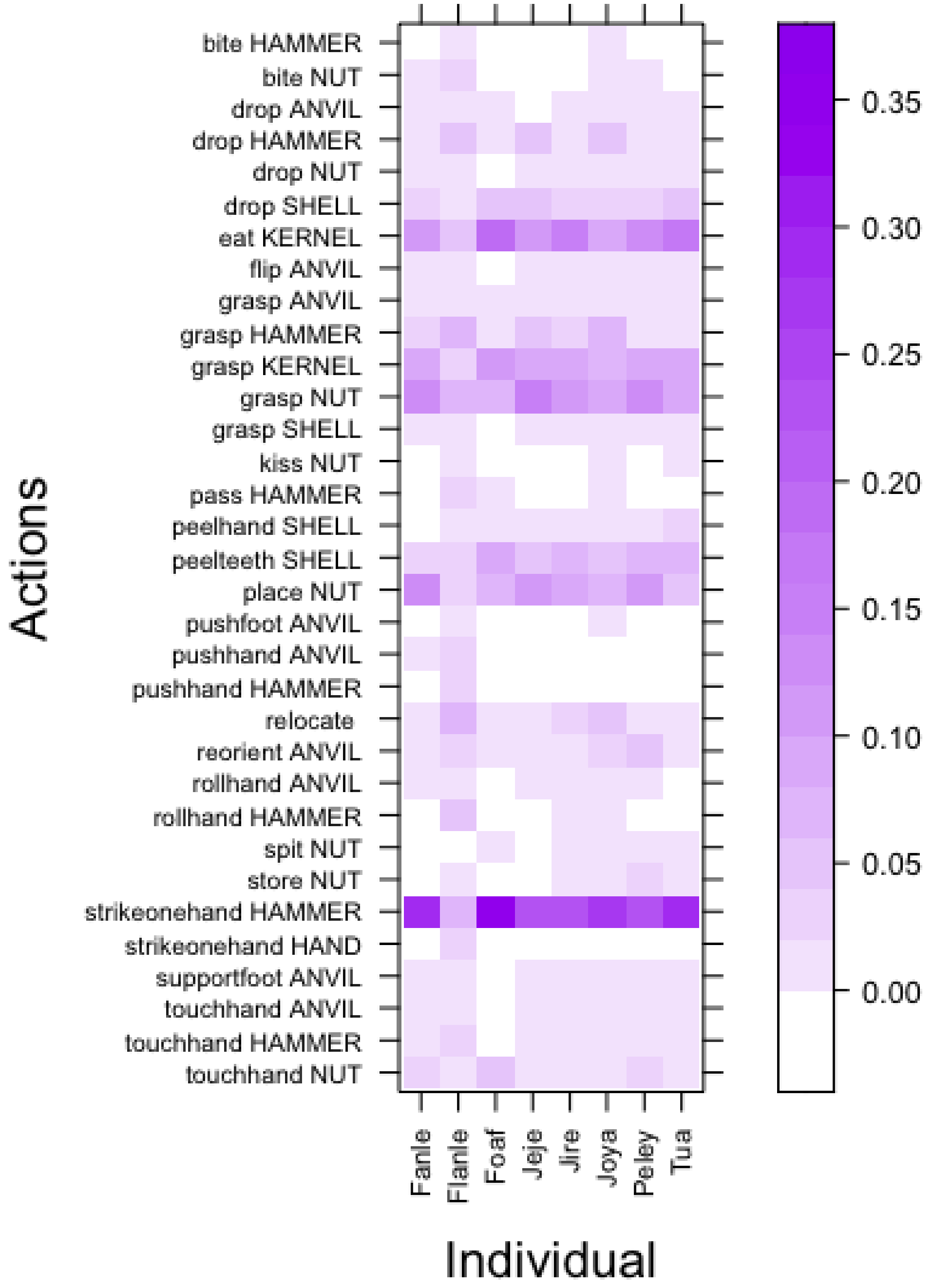


**Figure S2. Enlarged heatmap of action types used by chimpanzees during nut cracking.** The shading relates to the proportion of actions of a given action type that occurred in the data for each individual. This is a larger replication of figure 1.c. in the main manuscript.

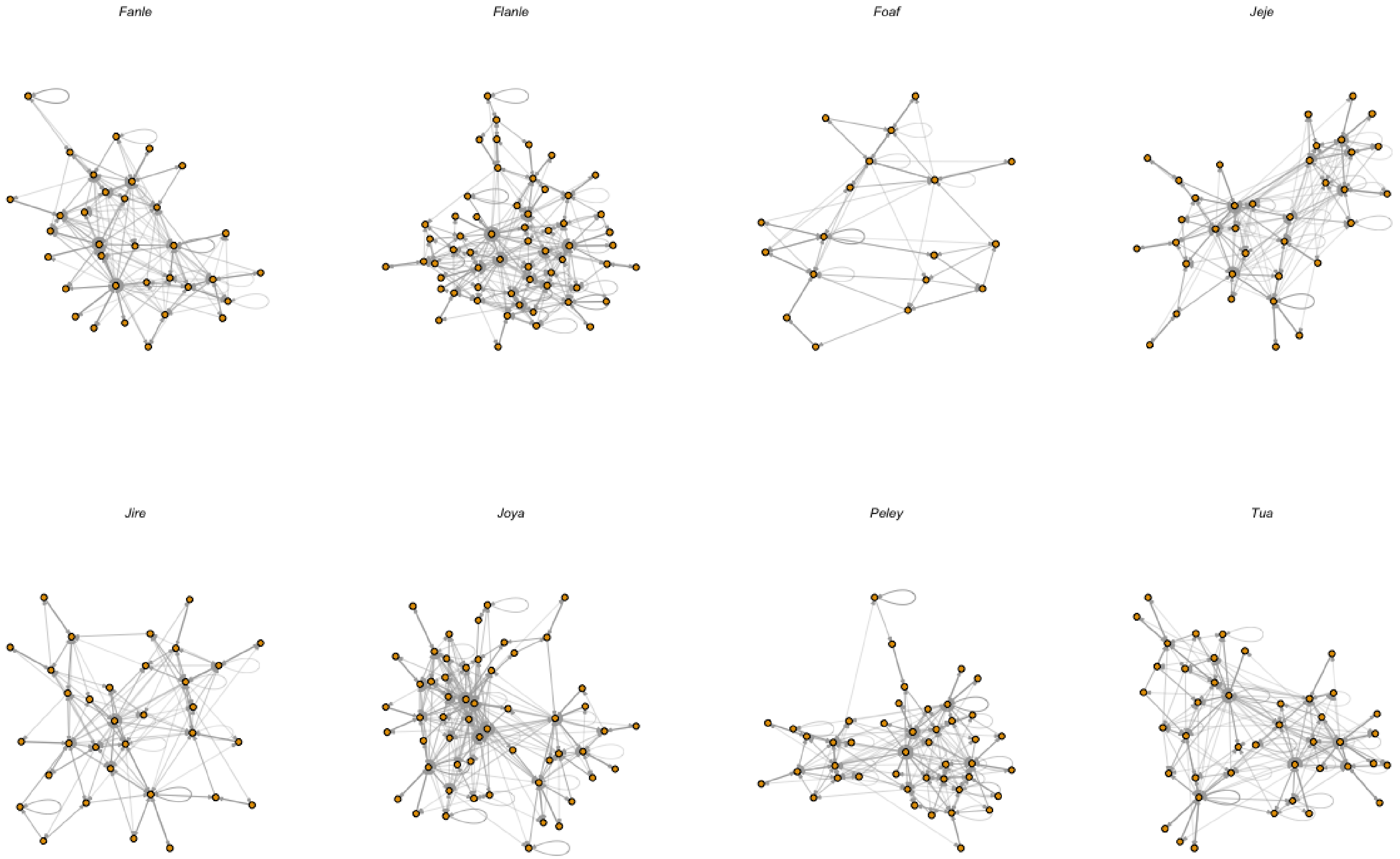


**Figure S3. Estimated Markov models for each individual.** Nodes in each network represent an action, e.g. *Drop Hammer, Spit Shell, Eat Kernel,* etc. Edges represent probabilistic transitions between the different action types. Probabilistic transitions are estimated from sequences using neighbouring elements – therefore, these models represent first order Markov chains, which contain no memory of states previously activated during sequence construction.

**Table S3. Maximum distances $MI_{Adj} > 0$ for sequences produced by chimpanzees and Markov models.** Confidence intervals were estimated using the exact method for the Poisson distribution. An asterisk (*) indicates that the maximum distance MIAdj > 0 in sequences recorded from chimpanzees (including sequences which feature repeating actions, and those that have been condensed) were greater than the 95% confidence interval for the mean maximum distance in Markovian Sequences. In these instances, we take this to mean that the distance was significantly greater for alpha = 0.05.

| Individual | Sequence Type | Max Distance MIAdj > 0 | Mean Max Distance MIAdj > 0 from Markov Data (Elements) | Variance in Max Distance MIAdj > 0 from Markov Data (Elements) | Lower 95% CI for Mean Max Distance MIAdj > 0 from Markov Data (Elements) | Upper 95% CI for Mean Max Distance MIAdj > 0 from Markov Data (Elements) | Individual Distance Outside of 95% CI for Markov? |
|---|---|---|---|---|---|---|---|
| Fanle | Observed | 16 | 9.39 | 15.1 | 5.06 | 16.3 | * |
| | Condensed | 15 | 7.93 | 7.30 | 4.07 | 14.3 | * |
| Flanle | Observed | 100 | 16.56 | 147 | 10.3 | 25.4 | * |
| | Condensed | 8 | 6.25 | 5.12 | 2.97 | 12.0 | |
| Foaf | Observed | 8 | 9.74 | 9.75 | 5.31 | 16.7 | |
| | Condensed | 9 | 8.18 | 10.07 | 4.24 | 14.7 | |
| Jeje | Observed | 21 | 11.06 | 20.1 | 6.24 | 18.5 | * |
| | Condensed | 10 | 8.4 | 5.33 | 4.39 | 15.0 | |
| Jire | Observed | 10 | 7.61 | 13.1 | 3.85 | 13.9 | |
| | Condensed | 8 | 6.81 | 6.68 | 3.33 | 12.8 | |
| Joya | Observed | 20 | 10.27 | 8.58 | 5.68 | 17.4 | * |
| | Condensed | 8 | 8.27 | 7.45 | 4.30 | 14.8 | |
| Peley | Observed | 41 | 11.44 | 11.6 | 6.51 | 19.0 | * |
| | Condensed | 25 | 9.4 | 5.98 | 5.07 | 16.3 | * |
| Tua | Observed | 19 | 11.57 | 9.02 | 6.61 | 19.1 | * |
| | Condensed | 15 | 8.12 | 5.60 | 4.20 | 14.6 | * |

**Table S4. AICc scores for $MI_{Adj}$ decay models (where data includes repeats of actions).** For each individual, Max Element Distance highlights the greatest element distance included in the $MI_{Adj}$ datasets, and models are given in the order of preference (as determined by AICc). K represents the number of parameters estimated in each model. AICc describes the AIC score corrected for smaller sample size (see section 4.6). ∆AICc represents the difference between a given model's AICc score and the optimal model's AICc score for each individual. AICc.Wt describes the proportion of the total predictive power of all three models contained in each model, and Cum.Wt describes the accumulation of predictive power in order of model preference. LL represents the log-likelihood of each model given the data - AICc is estimated using both K and LL.

| Individual | Max Element Distance | Model | K | AICc | ∆AICc | AICc.Wt | Cum.Wt | LL |
|---|---|---|---|---|---|---|---|---|
| *Fanle* | 16 | Power-Law | 3 | -80.8 | 0.0 | 0.92 | 0.92 | 44.4 |
| | | Composite | 5 | -75.9 | 5.0 | 0.08 | 1.00 | 45.9 |
| | | Exponential | 3 | -50.6 | 30.2 | 0.00 | 1.00 | 29.3 |
| *Flanle* | 100 | Composite | 3 | -387.4 | 0.0 | 1.00 | 1.00 | 199.0 |
| | | Power-Law | 3 | -348.2 | 39.3 | 0.00 | 1.00 | 177.2 |
| | | Exponential | 3 | -268.2 | 119.3 | 0.00 | 1.00 | 137.2 |
| *Foaf* | 8 | Exponential | 3 | -27.6 | 0.0 | 1.00 | 1.00 | 19.8 |
| | | Power-Law | 3 | -15.2 | 12.4 | 0.00 | 1.00 | 13.6 |
| | | Composite | 5 | 8.8 | 18.7 | 0.00 | 1.00 | 24.1 |
| *Jeje* | 21 | Composite | 5 | -105.6 | 0.0 | 1.00 | 1.00 | 59.8 |
| | | Exponential | 3 | -85.4 | 20.2 | 0.00 | 1.00 | 46.4 |
| | | Power-Law | 3 | -76.0 | 29.6 | 0.00 | 1.00 | 41.7 |
| *Jire* | 10 | Power-Law | 3 | -38.0 | 0.0 | 0.99 | 0.99 | 24.0 |
| | | Exponential | 3 | -26.6 | 11.4 | 0.00 | 1.00 | 18.3 |
| | | Composite | 5 | -26.1 | 12.0 | 0.00 | 1.00 | 25.5 |
| *Joya* | 20 | Composite | 5 | -101.0 | 0.0 | 0.99 | 0.99 | 57.6 |
| | | Power-Law | 3 | -92.6 | 8.4 | 0.01 | 1.00 | 50.0 |
| | | Exponential | 3 | -67.4 | 33.6 | 0.00 | 1.00 | 37.5 |
| *Peley* | 41 | Composite | 5 | -245.5 | 0.0 | 1.00 | 1.00 | 128.6 |
| | | Power-Law | 3 | -226.3 | 19.2 | 0.00 | 1.00 | 116.5 |
| | | Exponential | 3 | -142.9 | 102.6 | 0.00 | 1.00 | 74.8 |
| *Tua* | 19 | Composite | 5 | -109.2 | 0.0 | 1.00 | 1.00 | 61.9 |
| | | Exponential | 3 | -78.7 | 30.4 | 0.00 | 1.00 | 43.2 |
| | | Power-Law | 3 | -64.9 | 44.3 | 0.00 | 1.00 | 36.2 |

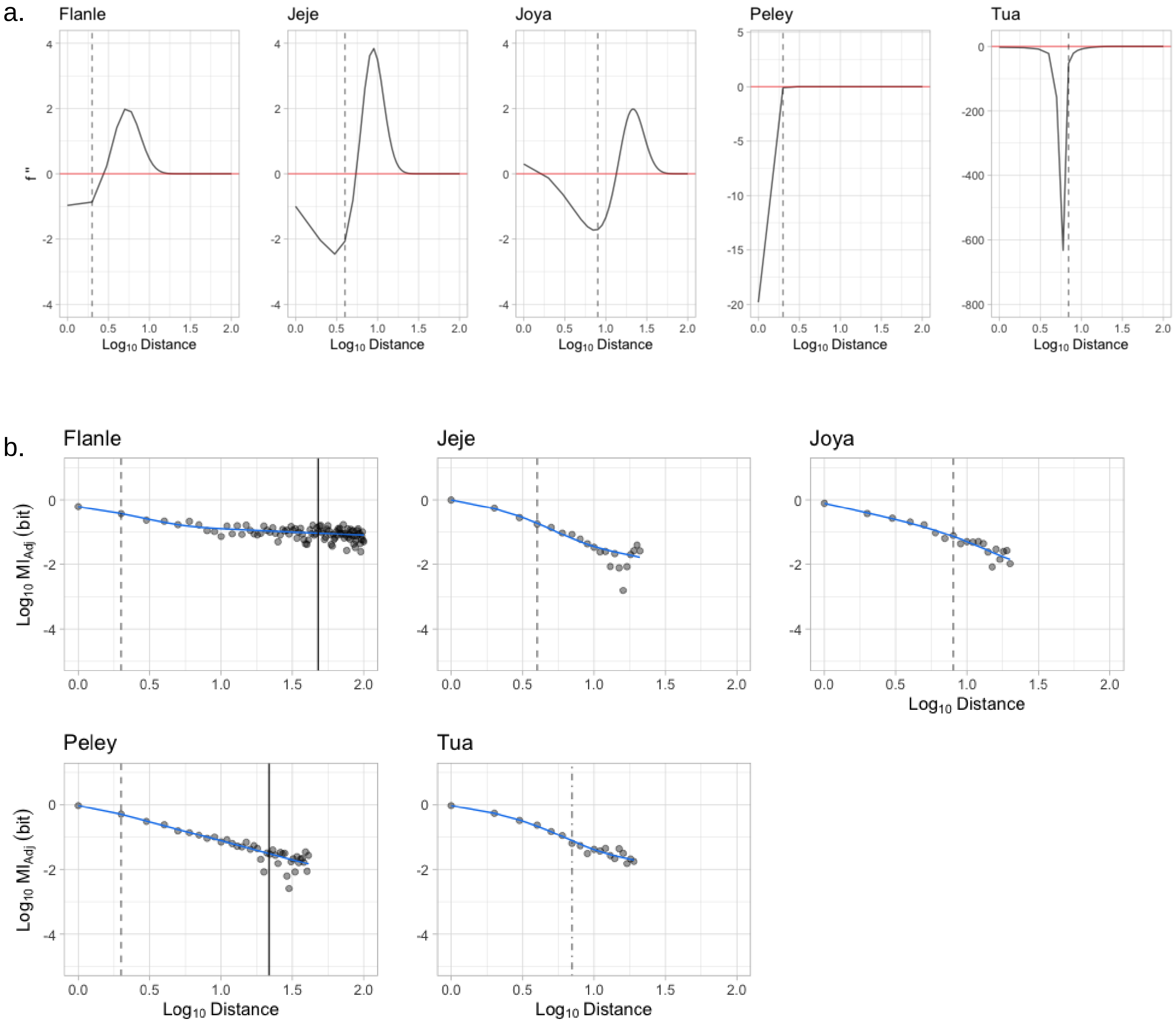


**Figure S4. Transition point profiles and composite decay models. (a)** Transition point profiles for each individual who showed a high preference for the composite model when data contained repeating elements. The point at which the profile is at its most negative identified where the change in gradient is also the most negative; the point immediately after this minimand indicates where the gradient is becoming less negative, and the composite model begins to 'pull upward' into a power-law decay. We mark this transition point on each profile with a dashed line. **(b)** The composite model for each individual for whom it offered the lowest AICc of all candidate decay models. On each composite model, the corresponding transition point is marked with a dashed line. For individuals whose average sequence length per nut is less than the maximum sequential distance $MI_{Adj} > 0$, the average sequence length per nut is marked with a solid black line.

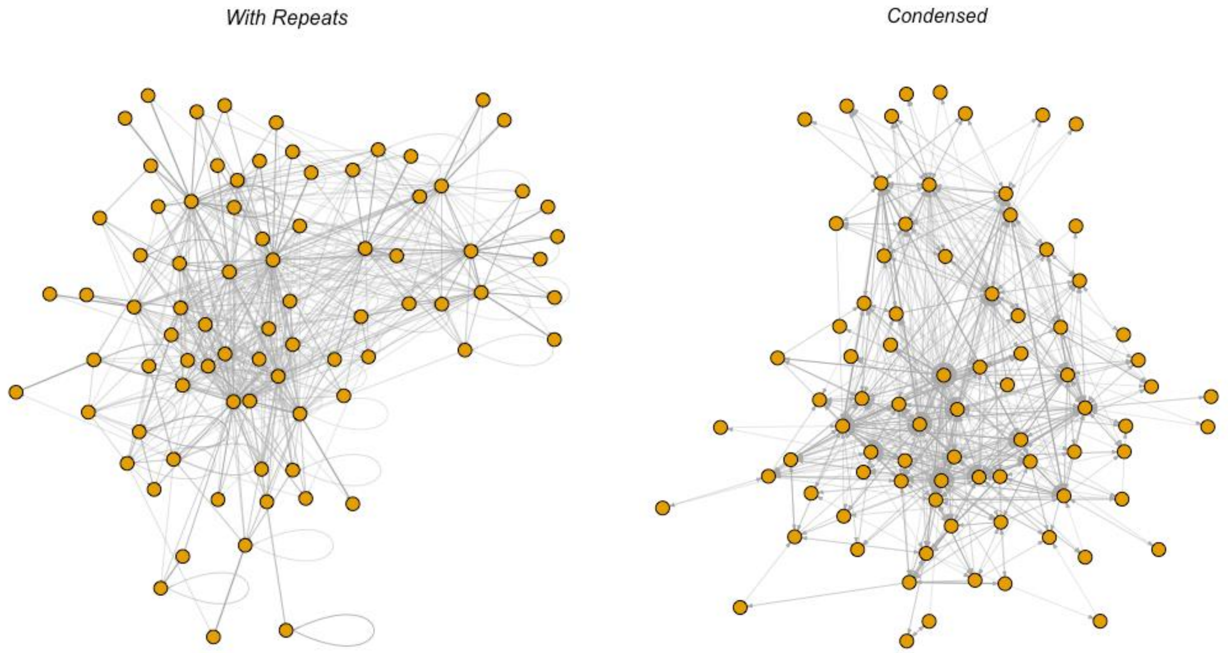


**Figure S5. Community Markov models.** The Markov models built using action-sequence data for all individuals; nodes represent action types, and edges represent the transition probabilities between action types in a sequence (darker = more likely). The Markov model on the left was built using action sequences with repeating actions. The model on the right was built using condensed data, where repeats were removed and replaced with a single action (e.g. the repeat 'Peel Shell', 'Peel Shell', would be replaced with a single codon: Peel Shell). Homogenized Markov models had identical connections to the graphs below, however outward transition probabilities from an action type were all equal; e.g. If there are two possible actions following 'Grasp Nut': 'Place Nut' (80% probability) or 'Drop Nut' (20% probability), these probabilities were replaced with an equal probability (50%).

**Table S5. Output of GLM investigating success of fitting all three decay models, as a function of the maximum distance over which MIAdj > 0.** GLMs are fit with binomial distributions, and Logit link functions. Distance relates to the maximum sequential distance over which $MI_{Adj}$ was greater than zero. *** means significant to $p < 0.001$; ** means significant to $p < 0.01$; * means significant to $p < 0.05$; . means significant to $p < 0.1$.

| Markov Model | Coefficients | Estimate | Std. Error | Z value | P value | Significance |
|---|---|---|---|---|---|---|
| With Repeats | Intercept | -9.2522 | 1.1292 | -8.193 | 2.54e-16 | *** |
| | Distance | 1.5266 | 0.1703 | 8.963 | < 2e-16 | *** |
| Condensed | Intercept | -1.5068 | 0.6556 | -2.298 | 0.0216 | * |
| | Distance | 0.5410 | 0.1071 | 5.053 | 4.36e-07 | *** |
